# A community-consensus reconstruction of Chinese Hamster metabolism enables structural systems biology analyses to decipher metabolic rewiring in lactate-free CHO cells

**DOI:** 10.1101/2025.04.10.647063

**Authors:** Pablo Di Giusto, Dong-Hyuk Choi, Athanasios Antonakoudis, Vikash Gokul Duraikannan, Pierrick Craveur, Nicholas Luke Cowie, Tejaswini Ganapathy, Kannan Ramesh, Santiago Benavidez-López, Camila A. Orellana, Natalia E. Jiménez, Leo Alexander Dworkin, James Morrissey, Igor Marin de Mas, Benjamin Strain, Norma A. Valdez-Cruz, Mauricio A. Trujillo-Roldán, Jannis Marzluf, Verónica S. Martínez, Leopold Zehetner, Claudia Altamirano, Ana Maria Vega-Letter, Bradley Priem, Haoyu Chris Cao, Martin Hold, Junyu Ma, Yi Fan Hong, Saratram Gopalakrishnan, Blaise Manga Enuh, Chaimaa Tarzi, Kuin Tian Pang, Claudio Angione, Jürgen Zanghellini, Cleo Kontoravdi, Hooman Hefzi, Michael J. Betenbaugh, Lars K. Nielsen, Meiyappan Lakshmanan, Dong-Yup Lee, Anne Richelle, Nathan E. Lewis

## Abstract

Genome-scale metabolic models (GEMs) are indispensable for studying and engineering cellular metabolism. Here, we present *i*CHO3K, a community-consensus, manually-curated reconstruction of the Chinese Hamster metabolic network. In addition to accounting for 11004 reactions associated with 3597 genes, *i*CHO3K includes 3489 protein structures and structural descriptors for >70% of its 7377 metabolites, enabling deeper exploration of the link between molecular structure and cellular metabolism. We used *i*CHO3K to contextualize transcriptomics and metabolomics data from a CHO cell line in which lactate secretion is abolished. We found the reduced glycolytic flux and enhanced TCA cycle flux were accompanied by an elevated NADH and PEP levels in these cells, consistent with experimental measurements. Leveraging *i*CHO3K’s structural annotations, we identified candidate binding interactions of NADH and PEP with glycolytic enzymes showing model-predicted differential flux, suggesting novel allosteric regulation associated with the observed decrease in glucose uptake and glycolysis. Overall, *i*CHO3K offers a valuable framework for systematic integration of omics data, improved flux predictions, and structure-guided insights, thus advancing CHO cell engineering and enhancing biomanufacturing efficiency.

## Introduction

Chinese Hamster Ovary (CHO) cells are the preeminent mammalian hosts for producing therapeutic proteins, due to their ability to make high titers of human-compatible therapeutic proteins^1,2^. Despite historical gains in their protein titers, further improvements in cellular efficiency remain a priority, particularly given rising demands for therapeutic protein^3–5^. Such demands are being addressed in part through advances in digital and automated platforms, such as bioprocess digital twins^6^, which are helping to reduce reliance on trial-and-error approaches by integrating real-time data, statistical analysis, and mechanistic modeling.

Within this context, genome-scale metabolic models (GEMs) are a key resource, offering insights into the metabolic reactions, cellular processes, and culture conditions critical to biomanufacturing processes^7–11^. However, the extent to which a GEM can accurately represent diverse metabolic states hinges on its comprehensiveness (i.e., its capacity to simulate all relevant metabolic reactions). The more complete and detailed a GEM is, the more likely it will capture a given cellular metabolic state. The community-curated GEM reconstruction of the Chinese Hamster, *i*CHO1766, was carried out through a collaborative effort^8^. This reconstruction served as a cornerstone resource to understand Chinese Hamster metabolism at the genome-scale, to identify metabolic bottlenecks in CHO cell culture for rational cell engineering designs and process-optimization strategies^12–20^. Since then, disparate updates to the Chinese Hamster metabolic reconstruction have been proposed^9,10,21^, along with other works that expanded cellular processes beyond metabolism to include the secretory pathway^22,23^. Nevertheless, despite these advancements, a comprehensive reconciliation of available CHO metabolic information has not been achieved, and subsequent efforts have largely focused on refining the initial model, rather than expanding its metabolic scope; this has left many potentially relevant pathways unexplored.

To address this gap, we present *i*CHO3K, a community-consensus reconstruction that consolidates all existing CHO metabolic models and integrates newly identified metabolic reactions, curated from the literature. By incorporating over 3000 additional reactions from various databases and recently published expansions, we substantially broadened the network’s scope and improved the accuracy of its reaction and metabolite annotations. A global curation effort involving over 40 researchers worldwide significantly enriched gene annotations, culminating in a final model containing 3597 genes, 11004 reactions, and 7377 compartment-specific metabolites, making it the most comprehensive mammalian metabolic reconstruction to date. Beyond expanding its metabolic scope, *i*CHO3K incorporates extensive kinetic data, such as turnover numbers and enzyme molecular weights, which greatly enhance flux prediction accuracy. Additionally, the integration of structural biology data, including 3D protein structures for 3489 enzymes and structural descriptors for more than 70% of its metabolites, establishes a deeper connection between molecular structure and cellular metabolism.

To demonstrate the utility of *i*CHO3K, herein we employed it to elucidate the metabolic rewiring in a CHO cell line that was genetically engineered to remove the growth-inhibiting Warburg effect. Recently, we showed that lactate secretion could be eliminated in CHO cells by simultaneously deleting lactate dehydrogenase and pyruvate dehydrogenase kinases, producing the “Zero Lactate” (CHO-ZeLa) phenotype^24^. Although CHO-ZeLa shows improved growth despite lacking aerobic glycolysis, the metabolic rewiring and underlying regulatory mechanisms remain unclear. Here, we leveraged *i*CHO3K to integrate transcriptomic and spent media metabolomics data from CHO-ZeLa and wild-type CHO-S cells, to construct cell line-specific models that recapitulate the distinct metabolic states. We applied enzyme-capacity flux balance analysis (ecFBA)^12^ to these context-specific models and found a reconfiguration of central carbon metabolism, aligning with experimentally measured shifts toward oxidative metabolism and away from glycolysis. We then harnessed *i*CHO3K’s structural annotations to investigate allosteric regulation by NADH and phosphoenolpyruvate (PEP), metabolites that exhibited significantly elevated intracellular levels in CHO-ZeLa cells compared to the parental CHO-S. Guided by our ecFBA analysis, which pinpointed a subset of early glycolytic enzymes exhibiting reduced flux in CHO-ZeLa, we evaluated the binding potential of these enzymes for NADH and PEP. Our analyses revealed that PEP binds to an allosteric pocket on the Chinese Hamster ATP-dependent 6-phosphofructokinase (PFK), an interaction not previously reported in animals. This mechanism provides further insight into the reduced glycolytic flux and lower glucose uptake observed in CHO-ZeLa cells. Overall, these findings illustrate how *i*CHO3K bridges constraint-based modeling and structural biology to uncover novel regulatory insights, offering a comprehensive framework for understanding metabolic bottlenecks in CHO cells.

## Results

### A New Community-driven Genome-Scale Metabolic Network Reconstruction of the Chinese Hamster

In recent years, Chinese Hamster Ovary (CHO) cell modeling has advanced considerably through the development of multiple genome-scale reconstructions^8–10,23^. These efforts have substantially improved our understanding of CHO cell metabolism, but they still exhibit limitations in reaction coverage, gene annotations, and structural data integration. Moreover, these reconstructions stem primarily from *i*CHO1766 and thus lack a comprehensive reconciliation of all metabolic reaction databases, leaving numerous potentially relevant reactions unmodeled.

To address these gaps, we reconciled the existing models to create a unified foundation for further expansion. We integrated the reactions, metabolites, and gene annotations from *i*CHO1766, *i*CHO2101, *i*CHO2291, and *i*CHO2441, ensuring consistency and removing redundancies. Building upon this consolidated model, we incorporated >3000 additional metabolic reactions from other reconstructions and databases, including the human Recon3D^25^. These additions were mapped to Chinese Hamster genes through homology, allowing us to considerably expand the metabolic network’s scope and annotation. We then refined the resulting reconstruction by identifying and removing duplicate reactions and metabolites (**Figure 1A**).

**Figure 1.**
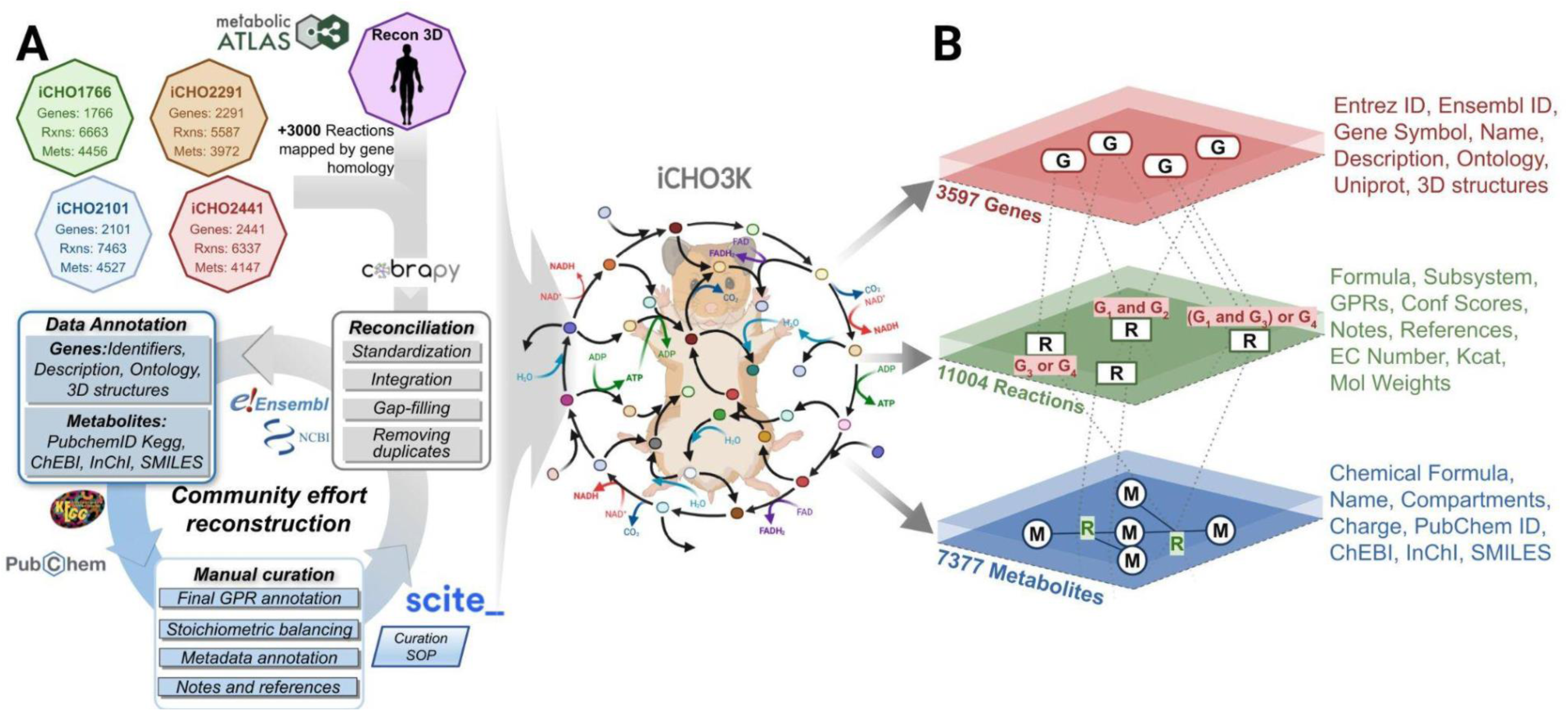
Schematic Overview of the *i*CHO3K Reconstruction Process: **(A)** *i*CHO1766^8^, *i*CHO2101^9^, *i*CHO2291^10^ and *i*CHO2441^23^ were reconciled and merged to define an initial set of reactions. Subsequently, >3000 reactions were mapped from Recon3D^25^ and other databases. This was followed by an annotation process, during which we retrieved various identifiers and attributes for genes and metabolites, including their 3D structures. The reconciled reconstruction, along with all its attributes and annotations, was then carefully curated manually through a community effort, following the guidelines established in the Manual Curation Standard Operating Procedure (See Materials and Methods). **(B)** The *i*CHO3K reconstruction integrates and expands upon existing models, incorporating 11,004 reactions, 7,377 metabolites and 3,597 genes with their different attributes annotated.

Gene-Protein-Reaction (GPR) annotations from the aforementioned reconstructions were included in the final draft. To enhance accuracy and completeness, we carefully curated the GPRs and metabolite annotations by reviewing their corresponding entries in the databases NCBI^26^, UniProt^27^, Rhea^28^, KEGG^29^, PubChem^30^, BiGG^31,32^ and ChEBI^33^, among others. Through a substantial community effort involving manual curation, we selected definitive GPRs for all reactions. This collaborative approach not only improved annotation quality but also yielded a shared resource for the CHO cell research community and biopharmaceutical industry.

Incorporating enzymatic information into metabolic models is increasingly recognized as essential for enhancing predictive accuracy^10^. Recent advancements in flux balance analysis have demonstrated that integrating kinetic parameters, such as enzyme turnover numbers (K_cat_) and molecular weights (MW), can narrow the solution space by accounting for the physical limitations of enzymes and thereby improve model reliability in simulating complex phenomena^34^. In our reconstruction, we provide comprehensive enzymatic annotations by mining the BRENDA database for these parameters (**See Materials and Methods**).

A novel feature of this reconstruction is the integration of structural data. We included 3489 3D protein structures, of which 2408 structures were generated using AlphaFold^35^ and Boltz-1^36^, and 1081 were retrieved from AlphaFold Protein Structure Database^37^. Furthermore, we integrated metabolite structures into this reconstruction by mapping InChI and SMILES strings from PubChem^30^ and other external databases, covering more than 70% of all metabolites in this reconstruction (**Supplementary Table 5**). This integration bridges the gap between metabolic network modeling and structural biology, enabling new avenues for analyzing enzyme functions, substrate specificities, and potential sites for metabolic engineering. To ensure transparency and reproducibility in the development of the reconstruction, every step of the process was meticulously recorded and thoroughly documented in a publicly available repository (https://github.com/LewisLabUCSD/Whole-Cell-Network-Reconstruction-for-CHO-cells).The resulting reconstruction, *i*CHO3K, represents a considerable advancement over previous models. It contains 3597 genes, 11004 reactions, and 7377 metabolites (**Figure 1B** and **Supplementary Tables 1-6**), making it the most comprehensive genome-scale mammalian metabolic reconstruction to date.

### *i*CHO3K considerably expands on reaction, gene and GPR coverage

To quantify the improvements of *i*CHO3K over its predecessors, we compared its content and function with previous CHO metabolic reconstructions: *i*CHO1766, *i*CHO2101 and *i*CHO2291. In this comparison, we excluded *i*CHO2441, as it shares the same metabolic reactions as *i*CHO2291, with the addition of the secretory pathway reactions. *i*CHO3K demonstrates a considerable expansion of the metabolic network, incorporating between 3500 - 4700 more reactions than earlier models. Moreover, it includes 1216 additional genes compared to the most extensive prior CHO reconstruction, *i*CHO2291 (**Figure 2A**).

**Figure 2.**
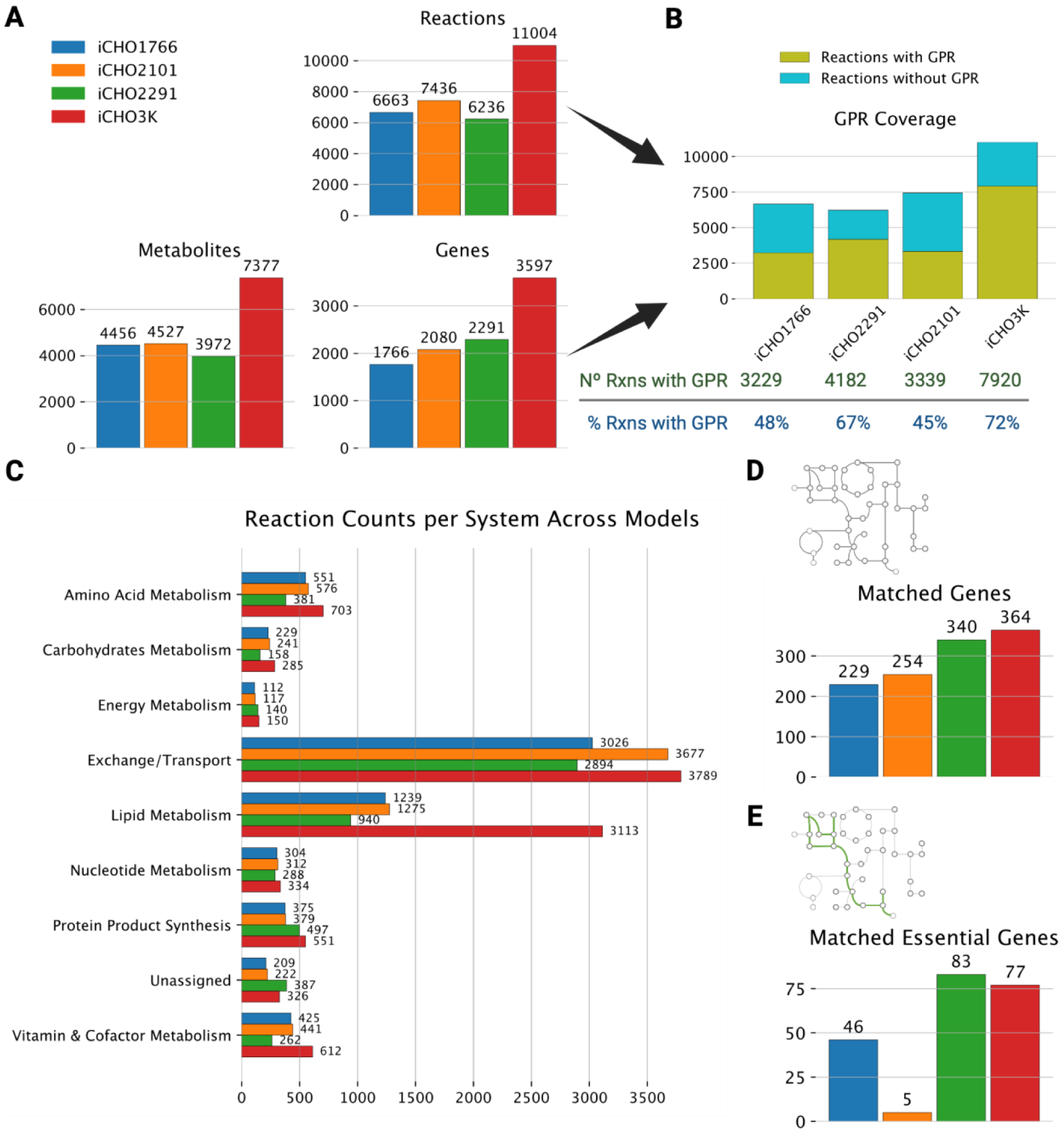
Comparison between *i*CHO3K and its predecessors: **(A)** Comparison of *i*CHO3K with previous CHO metabolic reconstructions (*i*CHO1766, *i*CHO2101, and *i*CHO2291) in terms of the number of reactions, metabolites and genes. **(B)** The proportion of reactions in *i*CHO3K with comprehensive gene-protein-reaction (GPR) annotations, covering 7,920 reactions (72% of the total), which considerably enhances transcriptomic and proteomic data integration from CHO cell lines. **(C)** Comparison of the number of reactions per system in each reconstruction, highlighting the expanded metabolic pathways in *i*CHO3K. **(D)** Experimentally validated essential genes included in each CHO reconstruction. **(E)** Correlation between experimentally validated essential genes and gene essentiality simulations, where the *in silico* deletion of the associated reactions reduces the biomass flux to less than 10% of its theoretical maximum.

A standout feature of *i*CHO3K is its comprehensive GPR annotation. The model includes 3,597 genes with detailed associations to 7,920 reactions, accounting for 72% of the total reactions in the reconstruction (**Figure 2B**). This level of annotation is unprecedented in CHO cell models and considerably surpasses previous reconstructions. This enhanced GPR coverage offers several advantages: it greatly improves data integration for model contextualization, thereby enhancing the predictive accuracy of the models and providing deeper biological insights into CHO cell metabolism.

*i*CHO3K benefits from an updated reaction subsystem organization. We adopted a framework^7^ that classifies the metabolic network into eight principal systems: Amino Acid Metabolism, Carbohydrate Metabolism, Energy Metabolism, Exchange/Transport, Lipid Metabolism, Nucleotide Metabolism, Protein Product Synthesis, and Vitamin & Cofactor Metabolism. These systems collectively cover 100 distinct subsystems, each linked to a corresponding metabolic pathway within the KEGG database to facilitate integration with this database and the future expansion of this network (**Supplementary Figure 1** and **Supplementary Table 6**).

To thoroughly compare the characteristics of reactions in each reconstruction, we standardized the subsystem classifications across all models, aligning them with the eight principal systems mentioned above. This alignment allowed us to directly compare the number of reactions present in each system between *i*CHO3K and its predecessors. Our analysis revealed that *i*CHO3K surpasses previous reconstructions in the number of reactions within every principal system, indicating a comprehensive expansion rather than isolated additions in specific pathways (**Figure 2C**). Notably, in the system of “Lipid Metabolism,” *i*CHO3K has more than double the number of reactions compared to previous reconstructions. This substantial increase is likely due to the incorporation of reactions mapped from human reconstructions by gene homology, where lipid metabolic pathways are more extensively developed than in CHO models^25,38,39^.

To evaluate the functional coverage of *i*CHO3K, we compared its gene annotations with a dataset of experimentally validated essential genes, identified through a genome-wide, virus-free CRISPR screen conducted on CHO cells. This screen targeted 15028 genes, and identified 1980 genes that significantly impacted cell growth^40^. Given that these genes are involved in diverse biological processes, it is expected that not all of them will be linked to metabolic reactions^40^. However, a comparison across all metabolic reconstructions revealed that *i*CHO3K contains the largest number of genes overlapping with this dataset (**Figure 2D**), which indicates that this reconstruction includes more genes fundamental for cell growth.

To further validate this reconstruction, we performed a gene essentiality analysis to compare genes predicted to be critical for cell survival *in silico* with those experimentally validated. This was achieved by simulating *in silico* deletions and evaluating their impact on biomass production. Genes whose associated reaction deletions reduced biomass flux to less than 10% of its theoretical maximum were classified as essential for growth. Interestingly, while *i*CHO3K showed the broadest coverage of essential genes, *i*CHO2291 performed better in accurately predicting essential genes that were experimentally validated (**Figure 2E**). This discrepancy may be attributed to the additional reactions in *i*CHO3K, which could render some previously essential reactions non-essential due to the presence of alternative pathways capable of producing essential metabolites required for growth.

### Contextualized *i*CHO3K models reveal distinct metabolic profiles in Zero-Lactate CHO cell lines

Even though *i*CHO3K represents the entirety of the metabolic network of the Chinese hamster, not every enzyme is expressed at all times and in every context. Gene expression varies across different cell types and tissues. Using a GEM as template, omics data can be overlaid to identify the active metabolic pathways and condition-specific network adaptations^41–44^. To harness this capability, we generated context-specific GEMs by integrating RNA-Seq and spent media metabolite concentration data, acquired over a 14-day fed-batch process from CHO-ZeLa and CHO-S (WT) cells. The CHO-ZeLa cells were engineered through the simultaneous deletion of lactate dehydrogenase and pyruvate dehydrogenase kinases to eliminate the Warburg effect^24^ (**Supplementary Figure 2**), resulting in a phenotype that exhibits no lactate secretion and a reduced glucose uptake rate while maintaining WT-like growth rates.

To generate context-specific GEMs, we began with *i*CHO3K as the base model. We pre-processed the RNA-Seq data using the Standep toolbox^43^ to calculate the ubiquity score for each reaction in each sample. Next, we constrained each model with experimentally-determined uptake and secretion rates and used the RMF mCADRE toolbox^44^ for model reduction (**see Materials and Methods**). Finally, we compared the network topology and functional capabilities of the generated models (**Figure 3A**). This pipeline resulted in 27 context-specific models, comprising 9 models from CHO-S (WT) cells and 18 models from CHO-ZeLa cells, spanning four culture time points with replicates for each condition.

**Figure 3.**
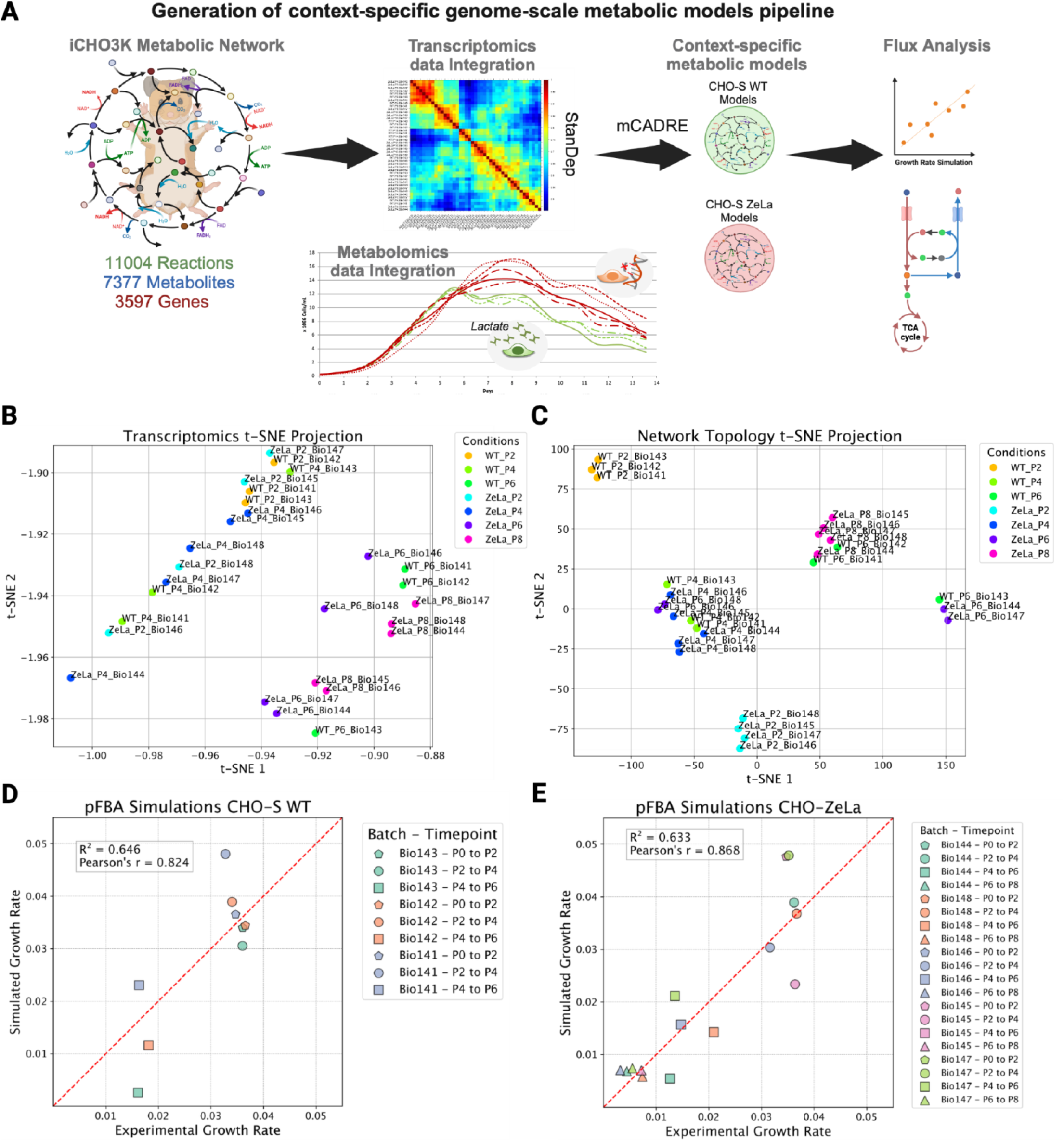
Distinct metabolic network structures and pathway activities revealed by contextualized modeling of Zero Lactate CHO cells: **(A)** Schematic representation of the context-specific genome-scale metabolic model reconstruction pipeline using *i*CHO3K. This process yielded 24 models representing wild-type (WT) and Zero Lactate (ZeLa) CHO cells at various culture days. **(B)** t-SNE embedding visualization of transcriptomic data from the 24 conditions analyzed. **(C)** t-SNE embedding visualization of the reaction structures of the 24 context-specific models, comparing models using the Hamming similarity method. Growth rate simulations using parsimonious flux balance analysis (pFBA) were compared against experimentally observed growth rates in CHO-S WT **(D)** and CHO-ZeLa (**E**) cells.

To evaluate whether the generated models captured the metabolic phenotypes of these cell lines, we performed t-SNE embedding comparison between the RNA-Seq data from the cell lines and the reaction structures of the models generated. Analysis of the transcriptomics data revealed a modest clustering among the conditions analyzed, particularly evident in WT_P2 and ZeLa_P8 models (**Figure 3B**). In contrast, the t-SNE plot of the reaction structures successfully clustered cell lines by their condition (WT or ZeLa) and culture time points (days 2, 4, 6, and 8) (**Figure 3C**). Interestingly, GEMs representing ZeLa_P4 (day 4) clustered together with WT_P4 models, suggesting that at this time point, there is minimal difference between the metabolic states of the two conditions. Additionally, some WT_P6 (day 6) models clustered with ZeLa_P8 (day 8) models, which correlates with experimental data indicating that WT cell lines reach the end of their exponential growth phase at day 6, while ZeLa cell lines do so at day 8 (**Supplementary Figure 3A**). This result highlights *i*CHO3K’s ability to reflect metabolic shifts corresponding to different time points and conditions in culture, and underscores the utility of integrating RNA-Seq and spent media data into *i*CHO3K to account for the topological and functional differences of metabolism, which is unclear in basic transcriptomic data.

*i*CHO3K structure allows us to compare both overall system coverage and detailed subsystem coverage, i.e., the relative number of reactions assigned to each system or subsystem across all generated models. While a system-level comparison provides a broad overview of inter-model differences, analyzing subsystem coverage offers more detailed insights within each system. We identified significant variation in practically all systems between WT and ZeLa models, particularly at days 4 (P4) and 6 (P6). Among the systems that varied the most, “Amino Acid Metabolism” stands out as the most differential between WT_P4 and ZeLa_P4, with the latter having more reactions in this system (**Supplementary Figure 4A**). A more detailed view revealed that “Tyrosine Metabolism” exhibits the greatest variation, contributing significantly to the differences observed in amino acid metabolism between WT and ZeLa cells at P4 (**Supplementary Figure 4B**). Tyrosine metabolism is directly linked to the tricarboxylic acid (TCA) cycle through its catabolic products, such as fumarate and acetoacetate, that serve as key substrates for energy production^45^. This connection suggests that the observed upregulation of tyrosine metabolism-related reactions in CHO-ZeLa cells may enhance the flow of these intermediates into the TCA cycle, thereby facilitating the metabolic adaptations necessary to eliminate the Warburg effect.

To simulate growth in the context-specific models generated, we employed parsimonious flux balance analysis (pFBA)^46^, maximizing biomass flux followed by minimization of the total flux sum. We then compared the predictions to observed growth rates. Our simulations revealed a stronger correlation between simulated and experimental growth rates in CHO-ZeLa cells (**Figure 3D**) than in WT cells (**Figure 3E**), indicating that the models and experimental constraints better capture the metabolic state of CHO-ZeLa cells than that of WT cells.

### ecFBA simulates experimentally observed phenotypes or metabolic fluxes

GEMs are valuable for simulating cellular metabolism, but their predictive accuracy depends on the constraints imposed during FBA. Traditional FBA approaches assume that reaction fluxes are only limited by stoichiometric constraints. However, recent advancements in FBA methodologies have introduced enzymatic capacity constraints (ecFBA)^10,12^, which account for the kinetic and physical limitations of enzymes catalyzing metabolic reactions. This approach reduces the feasible solution space based to the additional constraints from enzyme turnover numbers, i.e. K_cat_, and the total enzyme capacity of the cells.

Using the detailed enzyme annotation in *i*CHO3K, we employed ecFBA to explore the entire space of feasible metabolic flux distributions between the CHO-ZeLa and CHO-S cell lines under the given experimental constraints (**see Materials and Methods**), providing distributions for the fluxes through each reaction. We focused our analysis on models corresponding to day 4, when both cell lines are in exponential growth (**Supplementary Figure 3A**), to generate distributions for the fluxes through each reaction.

Our analysis revealed pronounced differences in the flux distributions of certain metabolic pathways between the two cell lines. In CHO-ZeLa cells, we observed a significant reduction in the early steps of glycolysis compared to CHO-S, a change that supports the diminished glucose uptake characteristic of the CHO-ZeLa phenotype^24^. Simultaneously, the simulations predict an increased flux toward the production of Acetyl-CoA (accoa) and L-alanine (ala) in CHO-ZeLa cells, suggesting a metabolic rerouting that favors these intermediates. Moreover, the enhanced flux through the TCA cycle in CHO- ZeLa cells aligns with their decreased reliance on aerobic glycolysis following the elimination of lactate secretion. Surprisingly, the flux distributions in the pentose phosphate pathway remain largely unchanged between the two models, indicating that the observed metabolic alterations are not affecting this pathway (**Figure 4A**).

**Figure 4.**
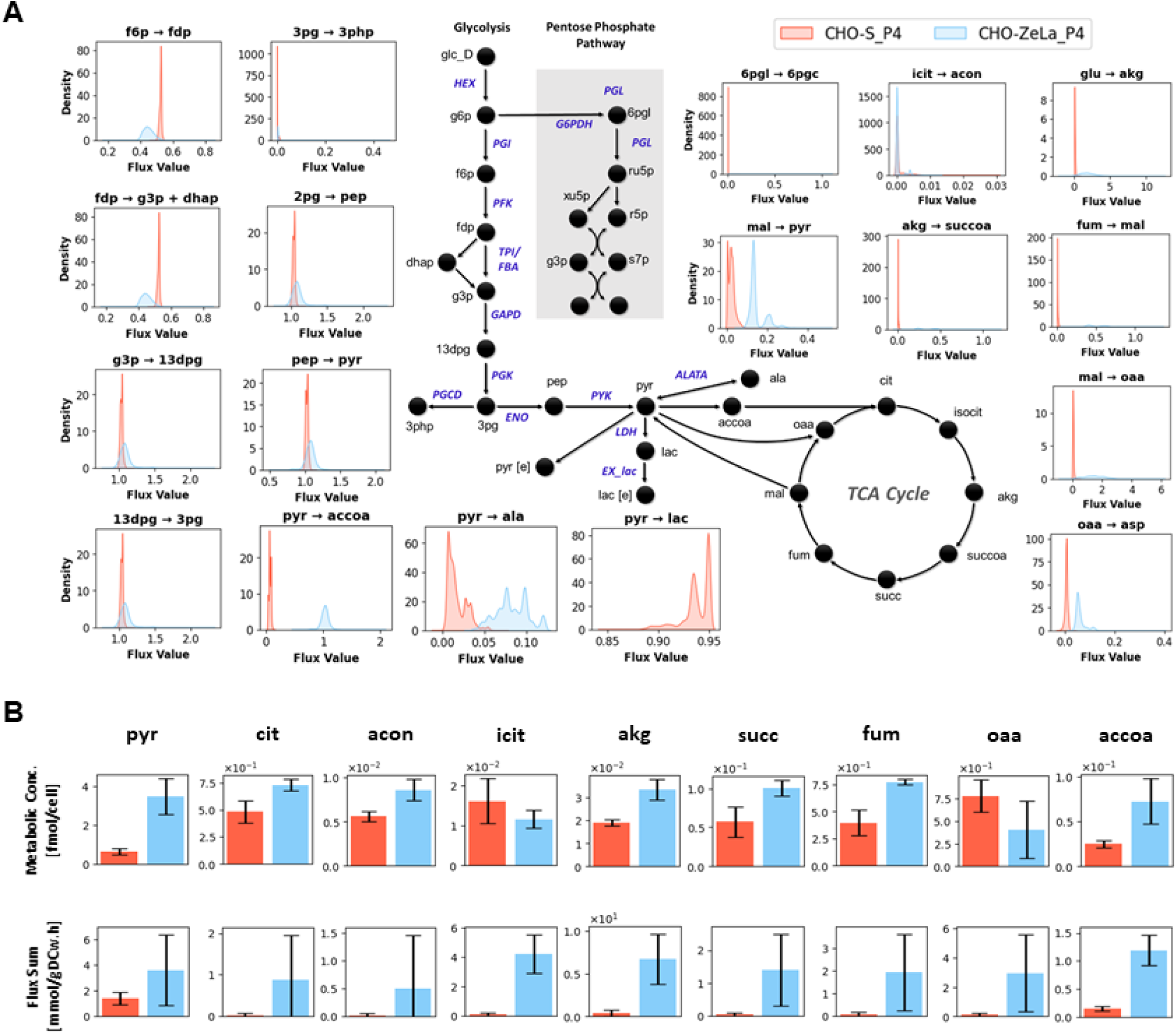
ecFBA reveals reduced glycolytic flux and elevated TCA cycle fluxes in CHO-ZeLa cells: **(A)** Flux distribution analysis between wild type (CHO-S) and CHO-ZeLa models at day 4 using the enzyme-constrained flux balance analysis (ecFBA). The results highlight distinct metabolic fluxes between the two cell lines in glycolysis, TCA cycle, and the pentose phosphate pathway. **(B)** Total metabolic flux through the TCA cycle in wild-type (CHO-S) and CHO-ZeLa cells at day 4 were compared to experimentally measured intracellular concentrations of TCA cycle metabolites. Bar charts display the predicted total flux through the TCA cycle for CHO-S and CHO-ZeLa cells as determined by ecFBA compared to the experimentally measured intracellular concentrations of the same TCA cycle metabolites. Orange bars represent wild-type (CHO-S) cells, while blue bars denote CHO-ZeLa cells.

Notably, while previous flux sampling analyses indicated a marked increase in flux toward PEP production, the current simulations show only a mild increase. This subtle change is consistent with a broader pattern observed for several metabolites under the altered metabolic state of CHO-ZeLa cells. Despite the modest flux increase predicted for PEP, experimental measurements during exponential growth revealed a significant accumulation of intracellular PEP in CHO-ZeLa cells (**Supplementary Table 7**). This discrepancy may be attributed to additional regulatory mechanisms, altered metabolite consumption rates, or substrate channeling effects that are not fully captured by the model.

To further validate our results, we compared the sum of fluxes through TCA cycle reactions predicted by ecFBA with experimentally-measured metabolite concentrations (**Figure 4B**). Overall, the simulations indicated an increase in TCA intermediates in CHO- ZeLa cells relative to CHO-S, confirming that ecFBA effectively captures the metabolic shifts observed experimentally. However, for certain metabolites: isocitrate (icit), and oxaloacetate (oaa), the model predictions deviated from the measured values. These discrepancies may be attributable to regulatory mechanisms not fully captured by the model, or inherent experimental variability in metabolite quantification.

These results demonstrate a clear metabolic reprogramming in CHO-ZeLa cells compared to wild-type CHO-S cells. Specifically, the simulations reveal a significant downregulation of early glycolytic flux in CHO-ZeLa cells, accompanied by an increased flux toward accoa, and the TCA cycle. This metabolic rerouting supports the observed reduction in glucose uptake and lactate secretion, hallmark features of the CHO-ZeLa phenotype. Moreover, the redirection of pyruvate away from lactate production toward acetyl-CoA formation for entry into the TCA cycle suggests that NADH levels may be elevated in these cells, a hypothesis that was confirmed experimentally (**Supplementary Table 7**).

### 3D structural analysis suggests inhibitory mechanisms CHO-ZeLa glycolysis

3D structure data greatly complements metabolic reconstructions, helping to bridge the gap between systems and structural biology. While GEMs are traditionally employed to predict and interpret metabolic fluxes and cellular phenotypes, the integration of structural data offers new opportunities to explore molecular mechanisms underpinning pathway-level functions^47^. Specifically, structural annotations provide insights into enzyme-substrate interactions, cofactor binding, and allosteric regulation, enabling a deeper understanding of enzyme function and specificity^35^.

ecFBA simulations and experimental measurements confirmed that the elimination of the Warburg effect in CHO-ZeLa cells decreased glycolytic flux, diminished cytosolic recycling of NADH, and increased PEP production. Consequently, the elevated levels of these two metabolites could inhibit the activity of glycolytic enzymes^48–52^, which would contribute to the significant reduction in glucose consumption in CHO-ZeLa cells^24^. To investigate this potential regulatory mechanism, we screened for possible off-target interactions of NADH and PEP with ten key enzymes from the early glycolytic pathway in CHO cells. The selection of these enzymes was guided by a dual-criteria approach. First, ecFBA of our CHO cell models revealed a consistent reduction in early glycolytic flux in the CHO-ZeLa phenotype compared to wild-type cells, suggesting that regulatory inhibition might be at play. Second, we prioritized enzymes known from the literature to interact with NADH or PEP as positive controls.

Currently, 176 CHO protein structures are available in the Protein Data Bank (PDB)^53^; however, none correspond directly to the ten glycolytic enzymes selected here for study. Thus, we relied on homologous structures from other species to identify PDB structures sharing the same fold as our enzymes of interest, and co-crystallized with NADH, PEP, or structurally similar molecules. This allowed us to assess the potential binding pockets on our CHO enzymes. Then, we used Boltz-1^36^ as a binding predictor to build the complexes of our enzymes with NADH or PEP, and compared against the binding sites identified in those homologous structures. These findings, summarized in

**Table 1**, indicate the likelihood of interactions between each enzyme and the two target metabolites (see **Supplementary Table 8** for additional structural parameters and confidence scores).

**Table 1:** List of enzymes analyzed and summary of the results: Set of enzymes from the early glycolytic pathway in CHO cells were analyzed for potential interactions with NADH and PEP, and results are summarized here. The symbols indicate the likelihood of these interactions: **`+++`**, enzymes with similar fold than target bind NADH or PEP; **+**, enzymes with similar fold than target bind ligand related to NADH or PEP; **`-`**, no structural evidence supports the potential binding of NADH or PEP to the enzyme.

|  | Uniprot ID | Enzyme Name | NADH | PEP |
| --- | --- | --- | --- | --- |
| 1 | G3H6T5 | *HEX* | + | - |
| 2 | P50309 | *PGI* | - | + |
| 3 | G3IDJ8 | *PFK* | + | +++ |
| 4 | G3I4H6 | *FBA* | - | + |
| 5 | P17244 | *GAPDH* | +++ | + |
| 6 | P50310 | *PGK* | + | + |
| 7 | G3HM99 | *PGAM* | - | + |
| 8 | A0A9J7GF98 | *ENO3* | - | +++ |
| 9 | A0A9J7GH21 | *ENO2* | - | +++ |
| 10 | A0A3L7IG89 | *PKI* | + | +++ |

As expected, our analysis revealed a high likelihood of interaction between NADH and GAPDH (P17244), which is consistent with NADH being the enzyme’s catalytic product. Similarly, high likelihood of interactions were observed between PEP and PKI (A0A3L7IG89), ENO3 (A0A9J7GF98), and ENO2 (A0A9J7GH21), aligning with the fact that PEP is the catalytic product of these enzymes. Interestingly, our results also indicate a highly probable interaction between PEP and PFK (G3IDJ8), despite this enzyme neither producing nor directly involving PEP in its reaction (**Table 1**).

PEP has been reported to inhibit PFK in other organisms^48,49^, with studies on homologous enzymes from *Bacillus stearothermophilus* proposing a mechanistic basis for this inhibition^54^. However, no such mechanistic explanation has been explored in the Chinese Hamster or other higher mammals. Our findings suggest that this regulatory mechanism is conserved in the Chinese Hamster, with PEP acting as a direct allosteric effector that modulates enzyme conformation and activity. This interaction could contribute to shifting glycolytic flux between bistable states, thereby fine-tuning metabolic output^52^. Such a regulatory role for PEP is particularly intriguing in the context of CHO- ZeLa cells, where the elimination of lactate production is accompanied by reduced glycolytic flux and enhanced oxidative metabolism, indicating that this off-target binding event may play a part in orchestrating the overall metabolic rewiring.

To further assess this mechanism, we applied the Boltz-1 binding predictor, followed by local minimization, to evaluate the PEP binding site in the Chinese Hamster PFK. The analysis suggested a binding site that mirrors the one observed in *B. stearothermophilus* (**Figure 5**). Notably, the *Bacillus* structures (PDB codes 4i4i and 4i7e) form a homo-tetramer composed of two interacting dimers, with each dimer binding two PEP molecules (**Figure 5A**). Structural alignment of all chains showed that, by symmetry, PEP occupies a single conserved binding site when examined at the monomer level (**Figure 5B**).

**Figure 5.**
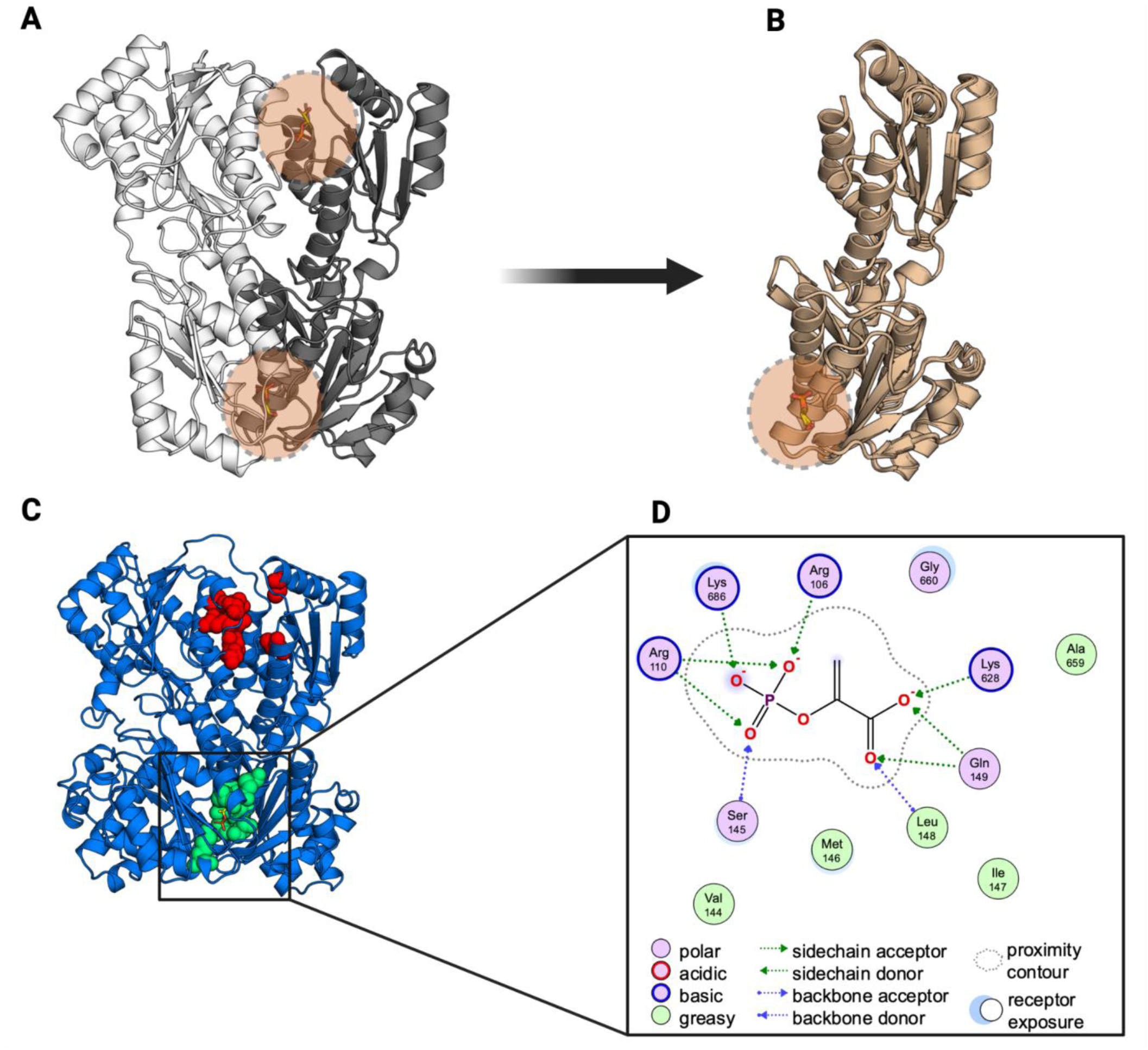
PEP binds symmetrically at a conserved inter-monomer interface in PFK: **(A)** Structures of *Bacillus stearothermophilus* PFK (PDB 4i4i and 4i7e; only 4i4i is shown) reveal PEP binds in different locations considering the dimeric states of the enzyme. **(B)** Structural alignment of all 16 chains from these PDB entries demonstrates that, by symmetry, PEP consistently occupies a single inter-monomer binding site. **(C)** A structural model of the CHO enzyme complexed with PEP predicts a similar binding site (green spheres) and, by symmetry, suggests a second site (red spheres). **(D)** In the predicted binding mode, PEP is stabilized by numerous hydrogen bond interactions with polar residues within the pocket.

Despite its longer sequence, the Chinese Hamster enzyme maintains a similar symmetric dimer fold, with PEP binding in a deeply buried pocket stabilized by hydrogen bonds from residues Arg-106, Arg-110, Ser-145, Leu-148, Gln-149, Lys-628, and Lys- 686 (**Figure 5C, green spheres; D**). Exploiting this symmetry, we propose a second equivalent binding site defined by residues Arg-491, Arg-495, Gly-528, Thr-531, Gly-532, Thr-264, and Thr-321 (**Figure 5C, red spheres**). Superposition of the Boltz-1 model with Bacillus X-ray structures (PDB 4i4i and 4i7e) reveals that while the phosphate group of PEP occupies the same position in the binding pocket, the carboxylic groups are oriented oppositely (**Supplementary Figure 5**).

Glycolysis can exhibit bistability, switching between low-flux and high-flux states, through intricate metabolite-mediated allosteric feedback and feed-forward controls^52^. Previous studies on rat and mouse PFK have shown PEP can be an inhibitor^48,49^, though the mechanism has not been explained. Our findings suggest a potential mechanism through which PEP could allosterically inhibit PFK, and contribute to the observed reduction in glycolytic flux following the knockout of LdhA and elimination of Warburg metabolism. Future studies employing high-resolution techniques such as X-ray crystallography or cryo-electron microscopy will be essential to confirm these interactions, and further metabolomics and fluxomics assays would be needed to fully characterize the molecular basis of the regulation of glycolysis in mammalian cells. However, this analysis provides an example of how detailed understanding into the metabolic rewiring of CHO cells can emerge from structural systems biology approaches, guided by metabolic flux simulations; thereby, supporting metabolic engineering and process optimization efforts.

## Discussion

Here we presented *i*CHO3K, the most comprehensive genome-scale metabolic reconstruction of the Chinese Hamster to date. By reconciling and integrating previous models^8–10^, we expanded the metabolic network to include 11004 reactions, 7377 metabolites, and 3597 genes. This includes expanded coverage across all metabolic systems, particularly lipid metabolism, enhancing our ability to predict cellular behavior under diverse conditions. Although the addition of alternative pathways sometimes led to discrepancies in essential gene predictions, this highlights opportunities for future refinements through additional regulatory constraints and pathway-specific adjustments.

Manual curation is critical for generating high-quality genome-scale metabolic reconstructions, as it ensures that each reaction, metabolite, and GPR association is evaluated and validated against experimental evidence and organism-specific data^55,56^. Through a community effort, we implemented a standardized protocol for manual curation that introduced a structured method for evaluating confidence scores and ensured integration with databases such as Rhea, BiGG, and KEGG. Engaging researchers from diverse laboratories worldwide, our collaborative process further resulted in a shared resource that benefits the entire CHO cell research community and biopharmaceutical industry. By fostering a culture of collaboration and data transparency, our approach creates a sustainable platform that will support ongoing updates and improvements.

The enriched annotations in *i*CHO3K enable more accurate predictions of CHO metabolism under diverse conditions, while also improving the model’s predictive performance. Notably, *i*CHO3K includes 7920 gene-associated reactions, which facilitated the integration of transcriptomics data for the generation of context-specific models of CHO-ZeLa and wild-type CHO-S cells; this revealed distinct metabolic adaptations underlying the zero-lactate phenotype^24^. Building on previous studies that underscore the importance of enzymatic constraints in phenomena like the Warburg effect^34^, we leveraged the extensive enzymatic annotation in *i*CHO3K to apply ecFBA for simulating intracellular metabolic fluxes. Our simulations predicted a reduced glycolytic flux and enhanced TCA cycle activity in CHO-ZeLa cells, aligning with prior experimental measurements^24^, and identified a subset of enzymes with notably lower turnover rates that may serve as potential targets for further mechanistic studies. Nonetheless, some discrepancies between simulated fluxes and experimental measurements for certain TCA cycle intermediates suggest that additional regulatory layers or methods may need to be incorporated to further refine model accuracy.

Integration of three-dimensional structural data with metabolic models has evolved over time, from early studies like the reconstruction of the *Thermotoga maritima* central metabolic network^47^ to more recent efforts, such as Recon3D^25^, which incorporated detailed protein and metabolite structural information on a genome-wide scale. Here, we have expanded the structural annotation in *i*CHO3K to cover 5427 compartment-specific metabolites and integrated high-quality 3D structures for 3041 enzymes. While Recon3D, primarily sourced protein structures from the Protein Data Bank and filled gaps with homology models, *i*CHO3K leverages state-of-the-art AlphaFold predictions to generate consistent, high-quality three-dimensional structure models for the majority of its enzymes. This approach overcomes limitations, such as unresolved residues and restricted isoform coverage inherent to experimental databases, and the intractable resource demand required to generate thousands of structures for a non-model organism. These enhanced structural annotations allowed us to identify potential regulatory interactions, such as the predicted allosteric inhibition of PFK by PEP. This provides new insights into the regulation of glycolysis in CHO-ZeLa cells, and could aid in process optimization and metabolic engineering for diverse CHO cell lines. By integrating three-dimensional structural data with constraint-based metabolic modeling, *i*CHO3K provides a robust framework for exploring molecular mechanisms of metabolic regulation, ultimately bridging structural biology with systems-level analysis and enhancing the interpretative capabilities of genome-scale reconstructions.

## Conclusion

*i*CHO3K represents a valuable resource in Chinese Hamster and CHO cell metabolic modeling that unifies diverse layers of biological data into a single, dynamic framework. Unlike traditional models that focus solely on reaction stoichiometry and gene associations, *i*CHO3K integrates refined enzymatic annotations with high-fidelity structural data, enabling a detailed exploration of cellular regulation that bridges molecular details with whole-cell physiology. This comprehensive resource not only empowers mechanistic investigations but also offers a modular, community-curated platform that can evolve with emerging data. The collaborative effort exemplifies the power of community engagement in scientific advancement, resulting in a valuable tool for optimizing bioprocesses and facilitating the development of more efficient therapeutic protein production strategies.

## Materials and Methods

### *i*CHO3K Metabolic Reconstruction

#### Reactions

The initial draft of the updated Chinese Hamster genome-scale metabolic network was obtained by integrating *i*CHO1766, *i*CHO2101, *i*CHO2291, and *i*CHO2441. Reaction formulas and metabolite compartments were standardized for correct integration. Duplicated metabolites and reactions were removed. Metadata and annotations from the three reconstructions were included in the final draft. GPR relationships for all reactions were extracted from each of the three CHO original reconstructions. Furthermore, GPR information from Recon3D^25^, the most complete mammalian metabolic reconstruction up to date, was mapped to our reconstruction by Human-to-Chinese Hamster ortholog identification which led to the addition of 3,718 extra reactions from Recon3D (**see Section 1, Reactions.ipynb, <u>GitHub Repository</u>**). GPR information acted as a measure of gene annotation robustness per reaction, thereby guiding the prioritization of manual curation efforts towards reactions with inconsistencies in their GPR annotation between reconstructions.

To identify duplicate reactions, we first generated a model from the reconstruction using the COBRApy python package^57^. Then, we generated an unfolded version of the stoichiometric matrix by taking into account the forward and reverse reactions ^57,58^. Finally, the unfolded stoichiometric matrix was used to create a correlation matrix and pairs of reactions with a value of 1, or 100% of correlation, were considered as duplicated (**see Section 2, Reactions.ipynb, <u>GitHub Repository</u>**).

For the boundary reactions, Extracellular Exchange reactions were automatically added for all the metabolites present in the extracellular space compartment. Intracellular Demand and Sink reactions were manually added during the manual curation of the reconstruction to ensure model feasibility and proper network connectivity, particularly for metabolites with known intracellular turnover but lacking explicit transport or conversion reactions in the existing reconstruction.

While refining our metabolic reconstruction, we adopted the subsystem classification framework as proposed by Richelle et al^7^. This framework allowed us to systematically organize the metabolic network into eight principal systems: Amino Acid Metabolism, Carbohydrates Metabolism, Energy Metabolism, Exchange/Transport, Lipid Metabolism, Nucleotide Metabolism, Protein Product Synthesis, and Vitamin & Cofactor Metabolism. These systems collectively cover 100 distinct subsystems, each meticulously linked to a corresponding metabolic pathway within the KEGG database^29^. The resulting draft of the reconstruction contains 11004 reactions, 7377 metabolites and 3597 genes

#### Metabolites

Metabolites were systematically extracted from the compiled list of candidate reactions. For each metabolite, detailed information such as names, chemical formulas, and charges were gathered from the respective reconstructions from which they originated (**see Section 1, Metabolites.ipynb, <u>GitHub Repository</u>**). Additionally, metabolite information was enhanced by integrating data from public databases. Missing metabolite names, chemical formulas, and charges were mapped from the BiGG database^31,32^. Subsequently, metabolite names and chemical formulas facilitated the retrieval of corresponding identifiers from the PubChem database^30^, which also yielded InChI and SMILES strings for each metabolite (**see Section 2, Metabolites.ipynb, <u>GitHub Repository</u>**).

Duplicated metabolites were identified and removed using a two-step process. Initially, metabolites were screened for similarities based on names, formulas, and PubChem IDs. Subsequently, molecular similarities were computed using the SMILES representations of the metabolites. This involved transforming SMILES strings into molecular objects via RDKit (https://www.rdkit.org) and calculating Tanimoto and Dice similarities between the fingerprints. A threshold of 0.99 (indicating 99% similarity) was set as the cut-off to define identical metabolites. This rigorous approach ensures a thorough and accurate identification of duplicate metabolites in our dataset (**see Section 3, Metabolites.ipynb, <u>GitHub Repository</u>**).

The final metabolites dataset comprises a total of 7,377 metabolites with their names and chemical formulas carefully annotated. Of these, 5,265 have an associated PubChem ID, 5,335 are linked with an InChI string, and 5,397 have an SMILES string.

#### Genes

A list of all the genes in the metabolic reconstruction was extracted in Entrez ID format from the model generated on COBRApy. We retrieved additional information, including gene symbols, names, and descriptions, from NCBI. This retrieval process was conducted using the Entrez submodule from the Biopython Python toolset^59^. To further enhance the accuracy of our data, we then mapped the Entrez IDs to Ensembl IDs for the genome assemblies of the Chinese hamster (CriGri-PICRH-1.0, GCF_003668045.3) and CHOK1GS (CHOK1GS_HDv1, GCA_900186095.1). This ensured the reliability and consistency of our gene information across different genomic databases. Finally, we mapped all Gene Ontology (GO) accession IDs related to each gene using the Ensembl’s REST API ^60^.

In addition to basic gene annotations, we extracted NCBI Transcript IDs and NCBI Protein IDs associated with each gene to enrich functional annotations. This information was obtained by parsing each NCBI Gene XML record to identify the corresponding mRNA (transcript) and protein products. Specifically, transcript IDs were retrieved based on their accession numbers, prioritizing those labeled with the ‘NM_’ prefix (RefSeq curated transcript) when available, and defaulting to the ‘XM_’ prefix (model-predicted transcript) otherwise. Corresponding protein IDs were extracted from the associated protein products of these transcripts. This retrieval process ensured that each gene entry was linked to its transcribed and translated products.

#### Biomass Reaction

To construct our biomass reactions, we differentiated between a recombinant-protein producing and a non-producing CHO cell line, mirroring the approach we detailed previously^8^. In delineating the biomass reactions, we categorized the main components into four distinct groups: DNA, Lipids, RNA, and Proteins. Each category was represented by a specific reaction that generated a representative metabolite, encapsulating all elements within that category. These representative metabolites were then incorporated into the biomass reactions. For metabolite coefficients within the two biomass reactions, mean values were derived from cell composition data, utilizing information from eight non-producing CHO cell lines for one reaction, and data from five recombinant-protein producing CHO cell lines for the other^61^. Importantly, these biomass reactions are generic and do not represent any specific CHO cell line. For our flux simulations, however, we employed a specific CHO-S biomass reaction to more accurately reflect the metabolic state of the cells used in our experimental studies.

#### Community-effort Manual Curation

Given the size of this project, we initiated a community-wide effort to manually curate the new Chinese Hamster reconstruction. We reached out to several labs globally, fostering a collaborative effort. In total, 38 individuals from various labs signed up to assist with the manual curation. The manual curation effort was guided by our newly developed Manual Curation Standard Operating Procedure (**Supplementary Data; Manual Curation SOP**) to ensure consistency and accuracy in the curation process. The protocol provided detailed instructions for the manual curation process, specifying the databases to use for data retrieval and the procedures to follow based on the available information. It also included information on the databases used for manual curation, detailing the types of data expected from each database and the querying requirements. To streamline the curation process, the reconstruction was divided into 47 packages, each containing approximately 220 reactions with similar characteristics. Curators selected packages based on their expertise. The aim of the curation process was to ensure the reconstruction was accurate and comprehensive, integrating GPR associations from various Chinese Hamster and human reconstructions.

To curate all K_cat_ values in the reconstruction, the following approach was implemented: For irreversible reactions, K_cat_ values were obtained from the BRENDA database^62^, using the EC numbers as inputs. The curation process followed this organism hierarchy: Cricetulus griseus (Chinese Hamster) > Rattus norvegicus = Mus musculus > Homo sapiens > Other. For reversible reactions, values were extracted from Human1^38^. If suitable values were not found in these sources, we referred to the supplementary data from Yeo et al., 2020^10^. For all curated reactions, the corresponding reference organism and the source were documented. Comments were included if K_cat_ values were unavailable for the same substrate or if no K_cat_ value was found for reversible reactions.

#### 3D Protein Structure Generation

Protein IDs from the Gene dataset (**Supplementary Table 4**) were used to retrieve the protein sequence from the NCBI database. For each sequence, an exact match (100% identity and query coverage) was sought within the AlphaFold Protein Structure Database (AFDB)^37^. The search was performed using the sequences.fasta file available from the EBI FTP server (https://ftp.ebi.ac.uk/pub/databases/alphafold/), and only entries whose FASTA headers contained “OS=Cricetulus griseus” were considered. When a match was found, the corresponding AFDB identifier was used to download the associated PDB structure. This procedure yielded 1081 protein models. For sequences without an exact match in AFDB, structural models were generated locally using AlphaFold-2^35^ with the parameters “--max_template_date=2023-04-25” and “-- db_preset=reduced_dbs,” resulting in 1955 additional models. In cases where the computational demand of AlphaFold-2 was prohibitive, Boltz-1^36^ (with default parameters) was employed as an alternative, generating models for 453 proteins. All local structure predictions (AlphaFold-2 and Boltz-1) were performed on a CentOS Linux 7 computing station equipped with two Nvidia RTX A5500 GPU cards. In total, 3489 protein structure models were obtained.

#### Model

Two versions of the iCHO3K model are provided. The comprehensive version, “*i*CHO3K,” encompasses the full manually curated network: every reaction, gene, and metabolite included in the final reconstruction. In contrast, the streamlined version, “*i*CHO3K_unblocked”, has been refined by removing blocked reactions, dead-end metabolites, and those reactions that led to spurious ATP and Glucose generation loops. This reduced model is designed to be used for simulations and model contextualization tools. Both versions of the model remain completely unconstrained by experimental data, meaning no uptake, secretion, or other boundary conditions based on empirical measurements have been applied.

### Context-specific model generation

To generate context-specific metabolic models for each experimental condition, we first created a reduced version of the network by removing blocked reactions and dead-end metabolites to ensure that all remaining reactions could carry flux. We then calculated ubiquity scores for each reaction using RNA-Seq datasets of each condition with the Standep tool^43^, to quantify the consistency of gene expression across different conditions and identify reactions that are active in a specific context. Next, we determined the uptake and secretion rates for each condition based on spent media data collected from a fed-batch process at various time points. Here, we processed the concentrations of amino acids, glucose, lactate, ammonia, and organic acids into specific uptake and secretion rates. The collected time profiles were transformed into specific growth rate SGR, 1/day), integrated viable cell density (IVCD, cell·day/mL), and specific consumption/production rates of metabolites (Qm, pmol/cell/day) for each phase (days 0-2, days 2-4, days 4-6, days 6-8, days 8-12, and days 12-14) according to equations (1) and (2).

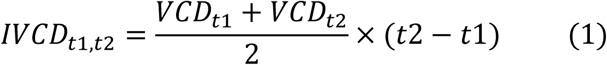

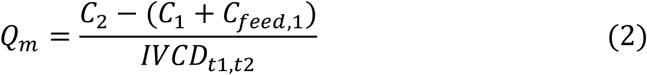

These rates reflect the consumption and production of metabolites by the cells under different experimental conditions. To convert the measured specific consumption/production rates (Qm) into mmol/gDCW/hr, we adopted a previously used conversion factor of 200 pg dry cell weight (DCW)^10^ per cell and calculated the averages and standard deviations from three replicate CHO-S WT batches and five replicate CHO-ZeLa batches. These values were then used to constrain the context-specific GEMs by applying boundary conditions based on the average value, adjusted by the standard deviations. Finally, we used both the ubiquity scores and the condition-specific uptake and secretion rates as inputs to the mCADRE tool^63^, setting the CHO-S biomass reaction (“biomass_cho_s”) as the objective function. Here, we defined protected reactions, which include biomass reaction, exchange reactions for nutrients and byproducts, as well as essential sink reactions, ensuring these key metabolic functions were retained in the context-specific models. The mCADRE algorithm integrates these data to refine the metabolic network, removing reactions unlikely to be active under the given conditions while preserving essential metabolic functions, thereby generating context-specific GEMs that better represent the observed cellular phenotypes.

### Metabolic network structure and transcriptomics data tSNE projections

An analysis of the metabolic network structures and subsystem coverage was conducted following a protocol similar to a previously reported method^38^. To compare the structures of the genome-scale metabolic models (GEMs), a binary reaction matrix was formulated, with rows representing individual reactions and columns corresponding to the GEMs being compared. A value of one in the matrix indicated the presence of a reaction in a GEM, while zero denoted its absence. To visualize the similarities and differences among the GEMs, t-distributed stochastic neighbor embedding (t-SNE) was applied to the binary reaction matrix, using the Hamming distance as the metric for measuring dissimilarity.

For the transcriptomics data analysis, genes were initially filtered to include only metabolic genes encompassed within iCHO3K, to ensure a direct comparison with the metabolic network structure. Gene expression values were normalized through a log₂(x + 1) transformation to normalize distributions, and minimize the impact of extreme values. Finally, t-SNE was applied to the normalized expression matrix using Euclidean distance as the dissimilarity metric, projecting the high-dimensional data into a two-dimensional space to facilitate visualization of gene expression patterns.

### System and subsystems coverage analysis

Variations in model structures were investigated by analyzing the coverage of metabolic systems and subsystems within the GEMs. System and subsystem coverage refers to the number of reactions present in a specific metabolic system or subsystem for a given model. To quantify system/subsystem coverage across different models, a comparison matrix was constructed. This was achieved by processing each GEM to count the number of reactions associated with each system or subsystem. To standardize the data a Z-score normalization was applied to each row of the comparison matrix. The normalized system/subsystem coverage data were visualized using clustered heatmaps. Hierarchical clustering was performed using the ‘average’ linkage methods and the ‘euclidean’ distance metric. Subsystems where all models exhibited zero coverage were excluded to enhance the interpretability of the heatmaps.

### Flux analysis

The contextualized *i*CHO3K model was adapted to represent the metabolic states of CHO-S (WT) and CHO-ZeLa conditions by integrating experimentally measured uptake and secretion. For each condition, these measured exchange rates were used as boundary constraints (see Context-specific model generation), and metabolites that were either not measured or not expected in the medium had their exchange flux bounds set to zero to prevent artificial nutrient uptake. To predict the context-specific growth rate, the CHO-S biomass reaction (“biomass_cho_s”) was set as the objective function, and parsimonious flux balance analysis (pFBA)^46^ was applied. pFBA minimizes the sum of absolute fluxes among all solutions and aligns with the biological principle of minimizing cellular machinery while maximizing growth.

Though pFBA has been an efficient simulation to predict growth rate, it falls short in fully capturing the intracellular metabolic behavior and translating the true enzymatic efficiency of the catalytic protein present with the cell. To better tackle this scenario, the available turnover number information from databases has been incorporated in the model to perform enzymatic capacity constraint FBA (ecFBA) simulation, which introduces an enzyme kinetic constraint (i.e., turnover numbers, 𝑘_𝑐𝑎𝑡_) in addition to the FBA constraints. By accounting for enzyme kinetics, ecFBA provides a more detailed and accurate representation of intracellular flux distributions. This makes it an ideal method for examining how cells manage their metabolic resources and adapt their intracellular metabolism. In this study, the ecFBA simulation minimizes the overall enzyme capacity factor while satisfying growth requirements, reflecting the cell’s optimization of resource use, formulated as follows^12^:

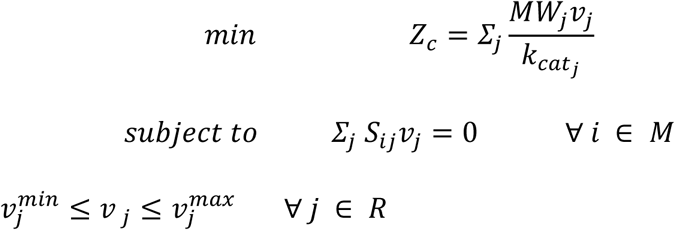

Where 𝑍_𝑐_ is the objective overall cellular enzyme capacity factor; 𝑣_𝑗_ is the flux solution of reaction 𝑗; 𝑆_𝑖𝑗_ is the stoichiometric coefficient of metabolite *i* in reaction *j*; 𝑣_*j*_^𝑚𝑖𝑛^ and 𝑣_*j*_^𝑚𝑎𝑥^ are the lower and upper bounds of the reaction 𝑗; 𝑀 and 𝑅 are the metabolite and reaction list respectively.

Deviations in the rate calculation due to experimental measurement errors are accounted for by constraining glucose uptake, lactate secretion and biomass reaction with ± 10% of measured rates as bounds. These constraints were found to be optimal to simulate the true physiology of the CHO-S and CHO-ZeLa cells while simultaneously minimizing the overall enzyme capacity factor. As we assume the cell is free to uptake required inorganics we leave the inorganics to freely exchange within the system, whereas all other measured metabolites’ uptake or secretion rate bounds span between 0 and experimentally determined flux value with 10% error (**Supplementary Figure 6**).

To complement reaction-based insights, we performed flux sum analysis, a metabolite-centric approach to examine compartmentalized metabolite turnover rates^64^. For rational comparison between the CHO-S and CHO-ZeLa cell lines, the flux solutions and the flux sums are normalized with respect to their individual glucose uptake flux and glucose flux sum value. The flux distributions of the glycolysis and TCA reactions on day 4 were compared between the cell lines. Whereas the Average flux sum values of the glycolytic and TCA metabolites were compared against the 13C metabolic flux analysis^24^.

Implementations of pFBA and ecFBA were conducted with cobrapy in a jupyter notebook and COBRA toolbox version 3.0^65^ in MATLAB 2020a. The Gurobi solver was used to solve the linear programming problems in pFBA and ecFBA^10^.

### Structural interaction analysis

To assess potential off-target interactions of NADH and PEP with the ten glycolysis enzymes of interest we applied the following strategy for each: First, we gathered available structural data on enzymes sharing the same EC number as the target proteins.

Protein structures of interest were identified using the Enzyme Structures Database (http://www.ebi.ac.uk/thornton-srv/databases/enzymes/) and the Advanced Search feature of the RCSB Protein Data Bank^53^, and the corresponding files were downloaded directly from the RCSB PDB. Next, we scrutinized this data to identify binding molecules that exhibit structural similarity to NADH and PEP, while also examining the binding contexts of these ligands within their respective enzymes, considering factors such as oligomeric status, the locations of catalytic pockets and binding sites, and the presence of any mutated positions.

Subsequently, we performed 3D structure superposition of the ligand-bound homologous enzymes with the AlphaFold2-generated model structures of the target enzymes^35^. This alignment was carried out using commands such as “align”, “super”, “cealign”, or “tmalign” in PyMOL version 2.6 (Schrödinger, LLC). Thereafter, we modeled the binding of NADH or PEP in the target enzymes using Boltz-1^36^, which allowed us to predict the binding complexes. Finally, the predicted binding configurations of NADH or PEP were compared with the observed binding pockets obtained from the superposition step, providing a comprehensive evaluation of potential off-target interactions.

To analyse the predicted binding of PEP in the ATP-dependent 6-phosphofructokinase from CHO, we used the model of the complex generated by Boltz-1 and apply a local minimization. This was done using Molecular Operating Environment (MOE) 2024.06 (Chemical Computing Group ULC, 910-1010 Sherbrooke St. W., Montreal) and its QuickPre*p* module with the standard settings, including Protonate3D to set the protonation states for pH 7.0, a threshold at 8Å to fix atoms away from the PEP ligand, an RMS gradient threshold of 0.1 kcal/mol/**Å²,** and Amber:EHT an all-atom force field combining EHT and Amber19. The 2D diagram of interaction was generated using the *Ligand Interactions* application.

## Acknowledgements

The authors would like to thank the following researchers who provided valuable feedback and/or an evaluation of a smaller number of reactions: Merle Rattay, Mareike Schulz, Tim Steffens, Shanti Pijeaud, Karl Gilmore, Dirk Mueller, Ali Safari, Swapnil Chaudhari, Nelson Ndahiro, and Caitlin Aamodt.

P.D.G. and N.E.L were supported by generous funding from Sartorius Stedim, the Georgia Research Alliance, and the National Institutes of Health (R35 GM119850). D.-H.C. and D.-Y.L. were supported by the National Research Foundation of Korea (NRF) grants (RS-2024-00341312 and RS-2024-00351458)) funded by the Korea government (MSIT). V.G.D and T.G. acknowledges the Half-Time Research Assistantship (HTRA) from the Ministry of Education, Government of India. M.L. acknowledges the funding support from the Indian Institute of Technology Madras for funding support through New Faculty Scheme (IP23242264BTNFSC009058). V.G.D., T.G., K.R. and M.L. also acknowledges infrastructure support from the Centre for Integrative Biology and Systems mEdicine (IBSE) and Robert Bosch Center for Data Science and Artificial Intelligence (RBCDSAI). N.A.V-C. and M.A.T.-R. were supported by Programa de Apoyo a Proyectos de Investigación e Innovación Tecnológica, Universidad Nacional Autónoma de México (PAPIIT-UNAM IN210822, IN218725, and IN211422) and Secretaría de Ciencia, Humanidades, Tecnología e Innovación (SECIHTI CF-2023-I-1549 and CF-2023-I-1248). N.A.V.-C. and M.A.T.-R. thank the sabbatical to PASPA-DGAPA-UNAM. N.L.C., I.M.d.M. and L.K.N were supported by The Novo Nordisk Foundation grants (NNF20CC0035580, NNF14OC0009473) and the GCHSP program (F-22010-24). C.A. was supported by the UK Biotechnology and Biological Sciences Research Council BBSRC (grant BB/Y01278X/1), and by Horizon Europe (grant 101182278). M.J.B. was supported by NSF Award #2114716. C.A.O. was supported by ANID Fondecyt de Iniciación en Investigación 11241251 and Open Seed Fund 2023 Escuela de Ingeniería UC. N.E.J. received funding from Núcleo MASH NCN2024037. J.M. and B.J. thank the U.K. Biotechnology and Biological Sciences Research Council (BBSRC) for their funding and support (BB/V509619/1, BB/V509620/1). C.Al. was supported by ANID Fondecyt Regular 1241453 and Fondef Idea ID23I10018. A.M.V-L was supported by Fondecyt Inicio 1220549.

## Author Contributions

P.D.G., D.-H.C., M.L., A.R., D.-Y.L. and N.E.L. conceived and designed the study and wrote the manuscript. P.D.G., D.-H.C, A.A., A.R. and N.E.L. led the network reconstruction and integration efforts. P.C. generated all the three-dimensional structures. P.D.G., D.-H.C, V.G.D. and P.C. created the figures and visualizations for the manuscript. P.D.G., D.-H.C, A.A., V.G.D., P.C., T.G., K.R. and M.L. performed the experiments and analyzed the data. N.L.C., L.K.N. and H.H. provided the experimental data. A.R., M.L. and N.E.L. reviewed and edited the manuscript.

Network curation contributions were provided by Santiago Benavidez-López, Camila A. Orellana, Natalia E. Jiménez, Leo Alexander Dworkin, James Morrissey, Igor Marin de Mas, Benjamin Strain, Norma A. Valdez-Cruz, Mauricio A. Trujillo-Roldán, Jannis Marzluf, Verónica S. Martínez, Leopold Zehetner, Claudia Altamirano, Ana Maria Vega-Letter, Bradley Priem, Haoyu Chris Cao, Martin Hold, Junyu Ma, Yi Fan Hong, Saratram Gopalakrishnan, Blaise Manga Enuh, Chaimaa Tarzi, Kuin Tian Pang, Claudio Angione, Jürgen Zanghellini, Cleo Kontoravdi, Michael J. Betenbaugh, Dong-Hyuk Choi, Vikash Gokul Duraikannan, Tejaswini Ganapathy, Pablo Di Giusto, Athanasios Antonakoudis and Anne Richelle.

All authors read and approved the manuscript.

## Declaration of generative AI and AI-assisted technologies in the writing process

During the preparation of this work the authors used ChatGPT-4 in a limited manner to improve the readability and language of the manuscript. After using this tool, the authors reviewed and edited the content. The authors take full responsibility for the content of the published article.

## Data and Code Availability

All data and code supporting the findings of this study are publicly available. The iCHO3K knowledgebase, models, datasets and scripts necessary to reproduce the analyses and figures presented in this manuscript are available in the GitHub repository at https://github.com/LewisLabUCSD/Whole-Cell-Network-Reconstruction-for-CHO-cells. Additionally, all the 3D structures generated for *i*CHO3K are hosted on Synapse.org and can be accessed at https://doi.org/10.7303/syn65473092.

## Supplementary Figures/Tables

**Supplementary Figure 1.**
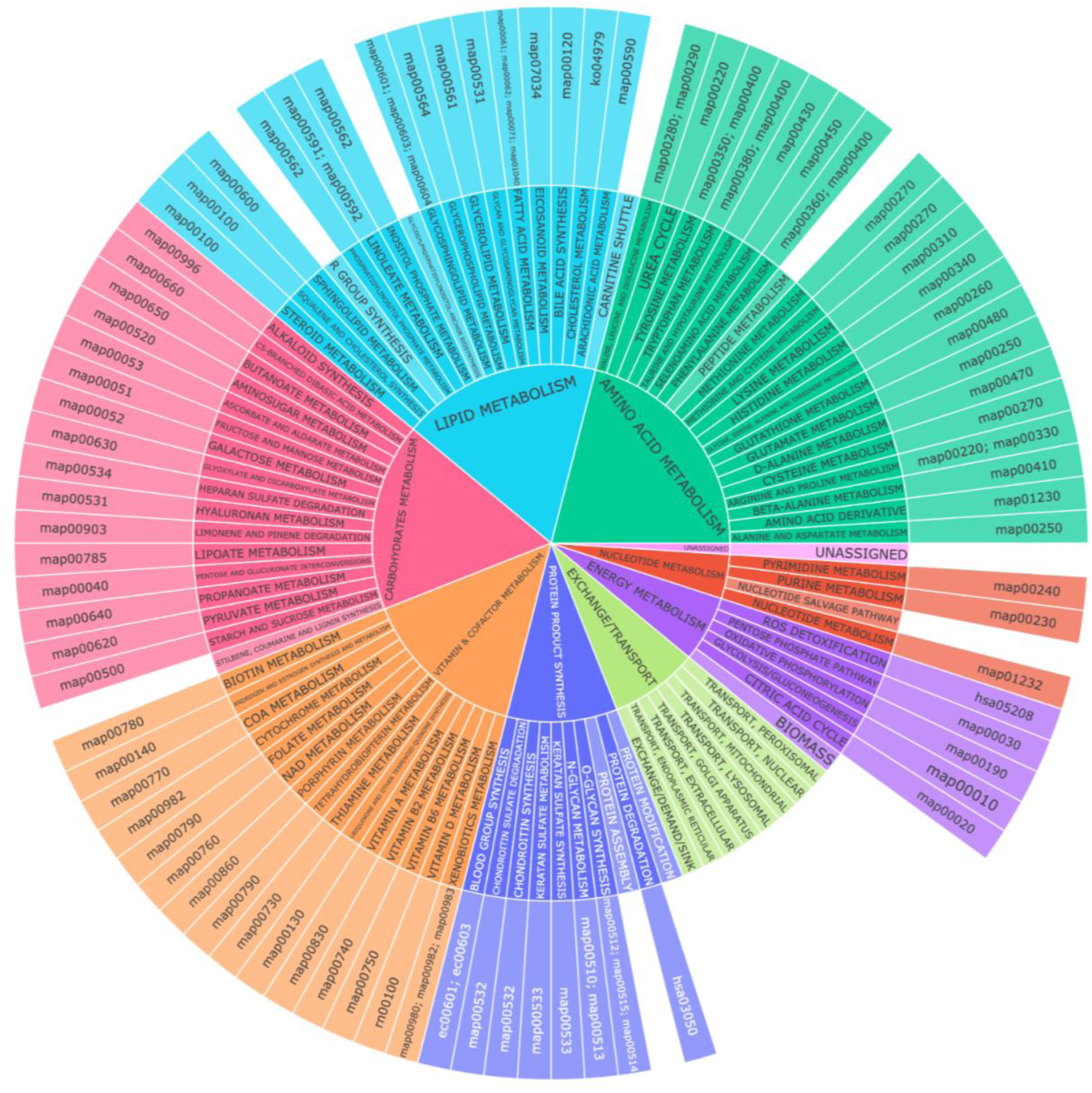
Systems and Subsystem Organization of *i*CHO3K: Subsystems corresponding to each reaction were redefined and organized into eight principal larger systems: Amino Acid Metabolism, Carbohydrate Metabolism, Energy Metabolism, Exchange/Transport, Lipid Metabolism, Nucleotide Metabolism, Protein Product Synthesis, and Vitamin & Cofactor Metabolism. Finally, each subsystem was mapped to an analogous KEGG pathway, allowing for pathway enrichment analysis.

**Supplementary Figure 2.**
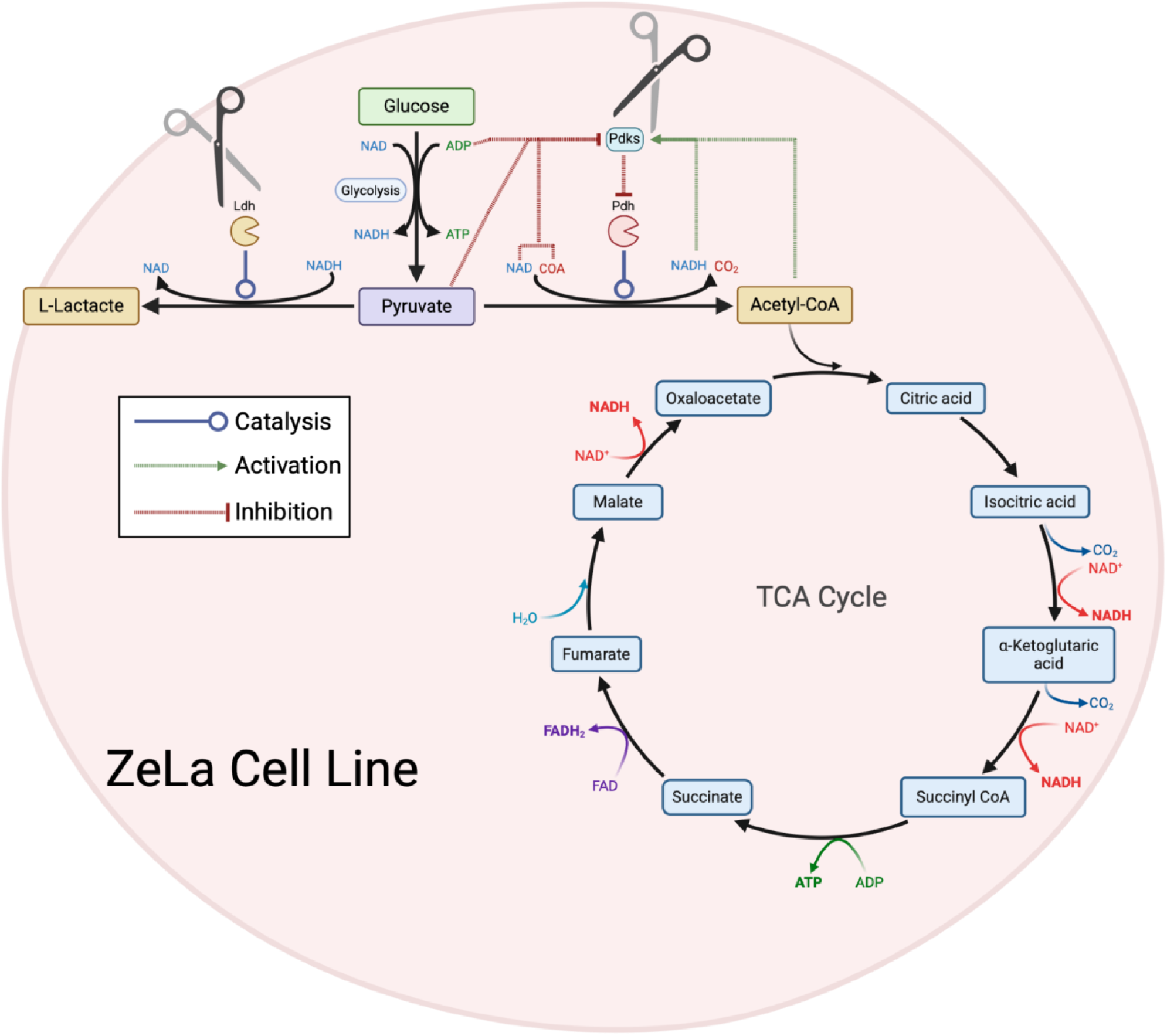
Schematic overview of ZeLa cell line: Pyruvate acts as a critical branch point between fermentation via lactate dehydrogenase (Ldh) and oxidative metabolism initiated by the pyruvate dehydrogenase (Pdh) complex. Pyruvate dehydrogenase kinase isoforms (Pdks) are regulated by the products and substrates of the Pdh reaction, forming a negative feedback loop that enhances lactate secretion when glycolytic flux is elevated. The Warburg effect is characterized by a high flux from glucose to lactate, with minimal conversion of pyruvate into the TCA cycle. The ZeLa cell line has been genetically modified to disrupt this feedback loop, thereby eliminating the Warburg effect.

**Supplementary Figure 3.**
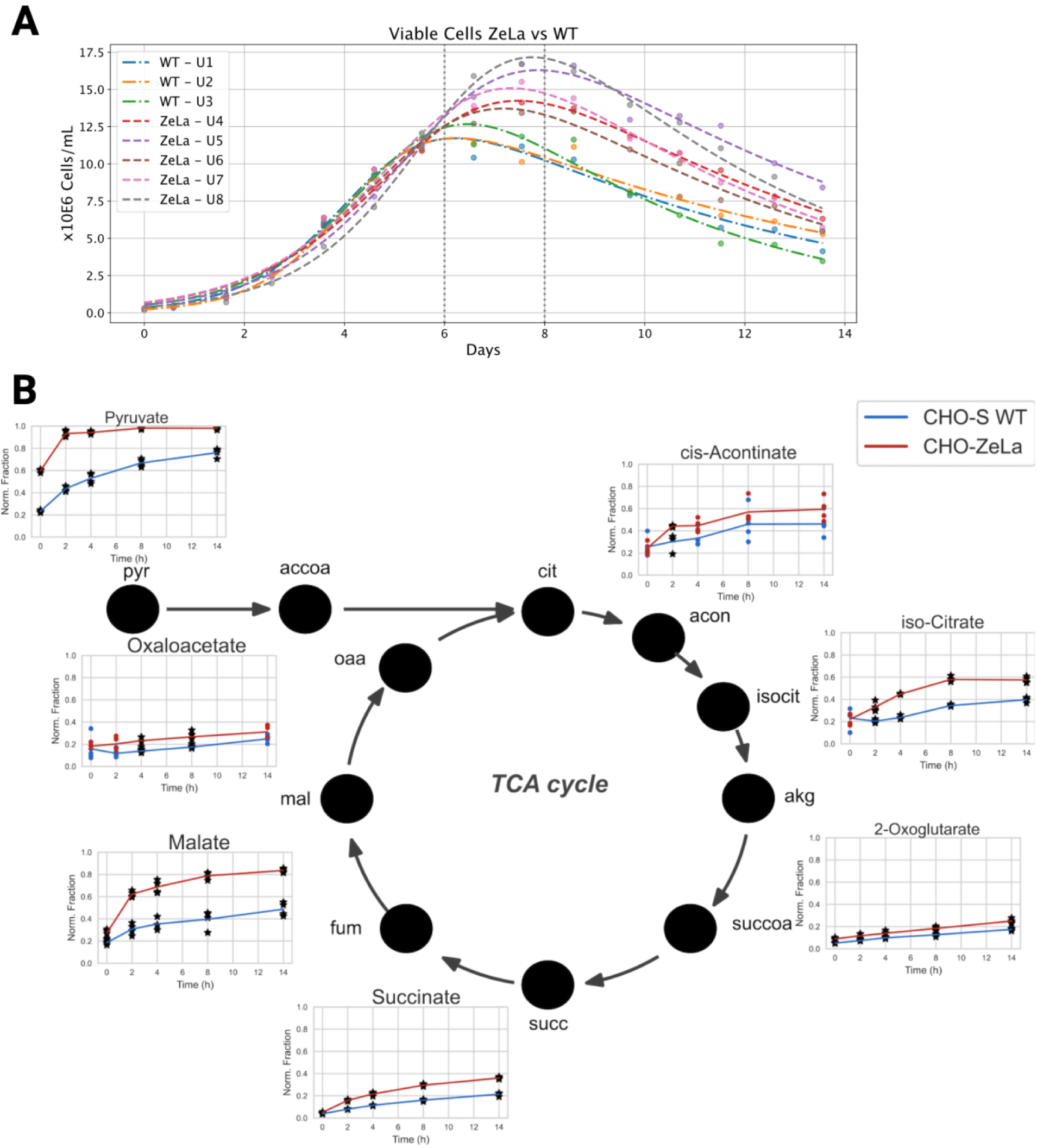
ZeLa cell lines exhibit increased maximum viable cell density (VCD) and enhanced oxidative metabolism: **(A)** VCD profiles of ZeLa and wild-type (WT) cell lines during a 14-day fed-batch cultivation process. Vertical dotted lines mark key cultivation times at days 6 and 8 where the higher VCD is observed for WT and ZeLa, respectively. VCD was calculated based on viable cell counts and adjusted culture volumes. **(B)** 13C labeling of TCA cycle metabolites following the addition of 13C labeled glucose during the mid-exponential phase, measured over time using liquid chromatography-mass spectrometry (LC-MS). Abbreviations-pyr: pyruvate, accoa: acetyl-CoA, cit: citrate, macon: aconitate, icit: isocitrate, akg: alpha-ketoglutarate/2-oxoglutarate, succcoa: succinyl-CoA, succ: succinate, fum: fumarate, mal: malate, oaa: oxaloacetate.

**Supplementary Figure 4.**
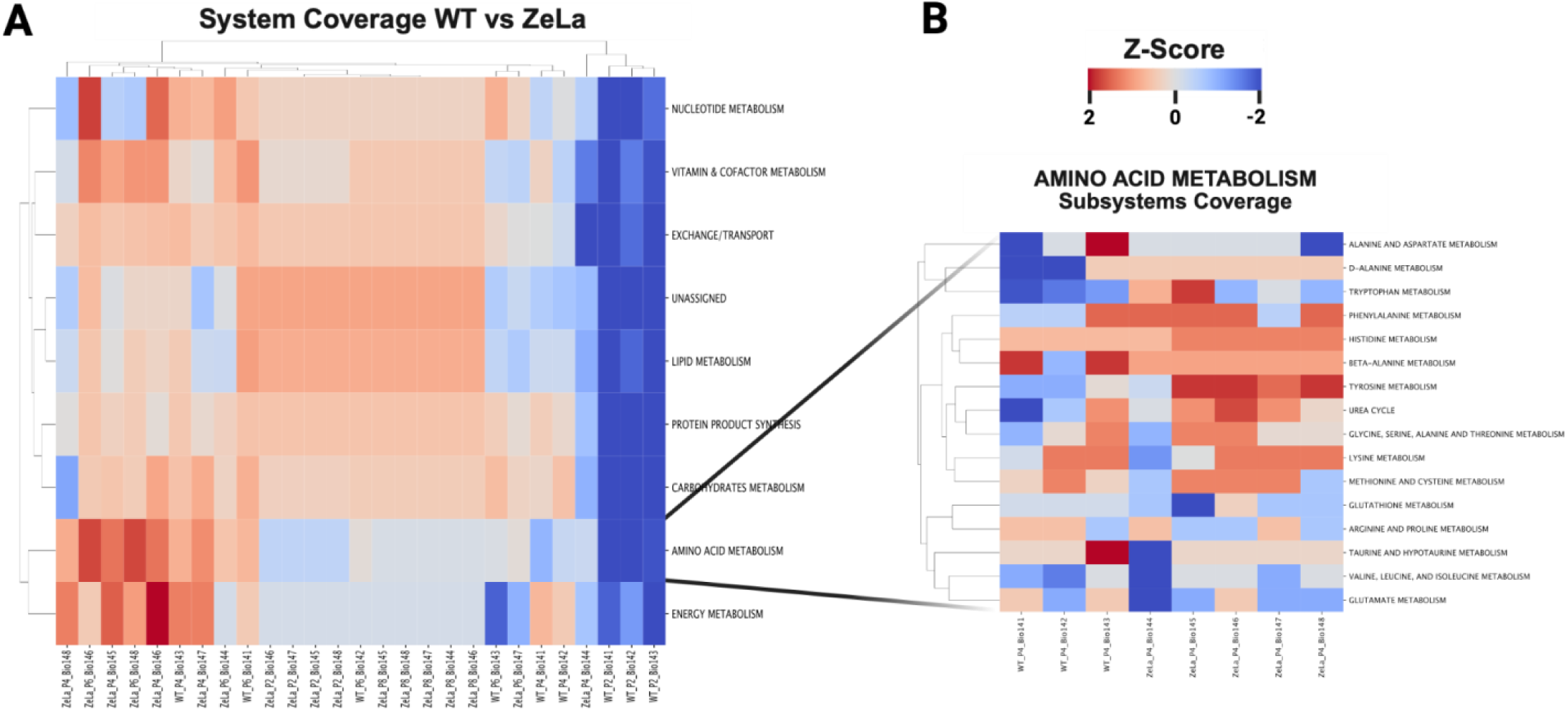
Comparison of system and subsystem coverage in context-specific GEMs: **(A)** Clustered heatmap showing Z-score normalized reaction counts for major metabolic systems across CHO-ZeLa and wild-type CHO-S models at days 2, 4, 6, and 8. Each row represents a metabolic system, while columns correspond to individual context-specific models. **(B)** A detailed breakdown of the Amino Acid Metabolism subsystem. Rows denote the specific reaction subsystems within the Amino Acid Metabolism system, while columns represent the corresponding models from both cell lines at day 4. The color scale ranges from blue (indicating below-average reaction counts) to red (indicating above-average reaction counts), and dendrograms illustrate the clustering of similar profiles.

**Supplementary Figure 5.**
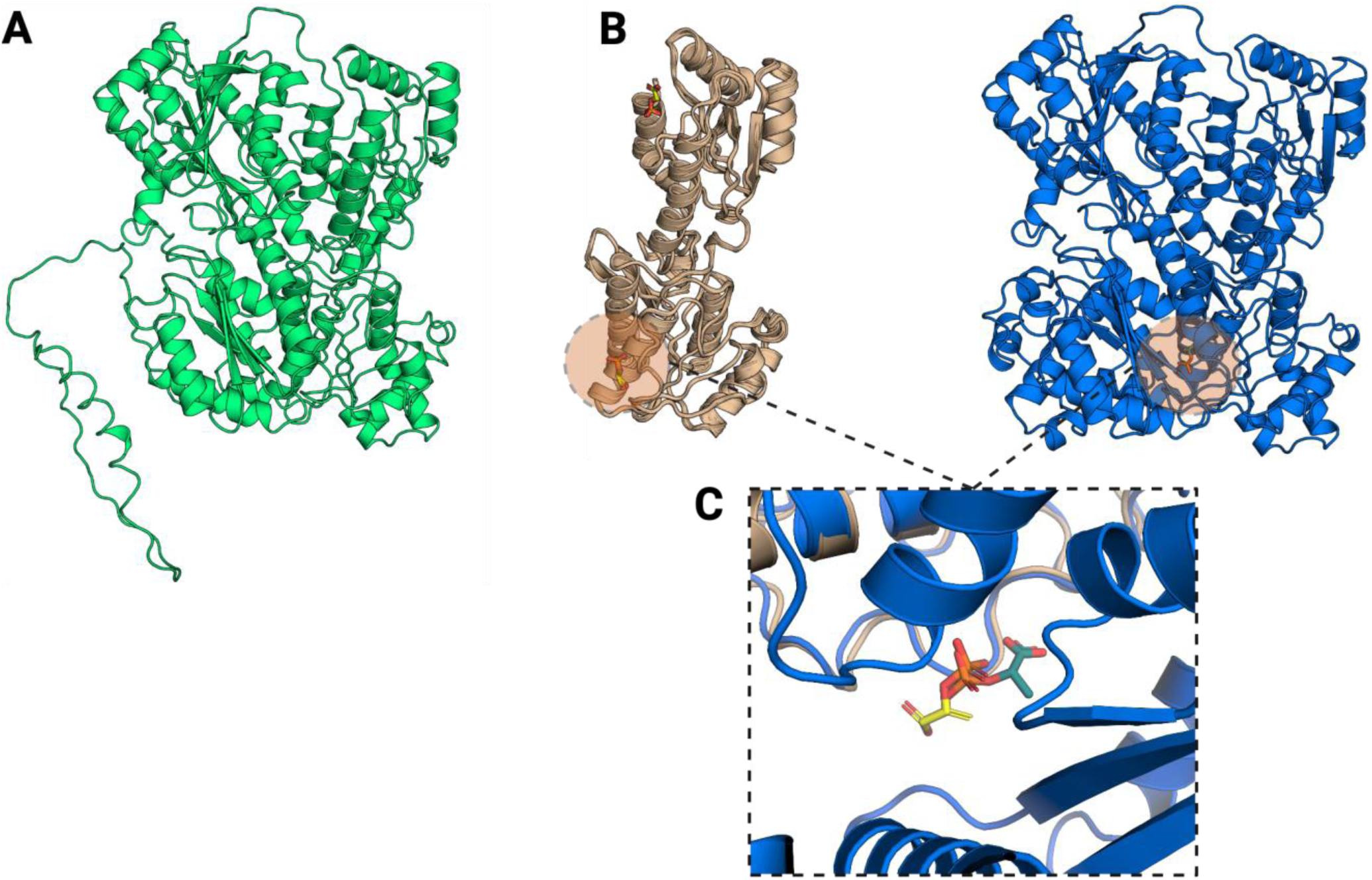
Carboxylic groups of PEP are oriented in opposite directions in the binding pocket of *B. stearothermophilus* and Chinese Hamster PFKs: (A) AlphaFold2-derived model of Chinese Hamster ATP-dependent 6-phosphofructokinase (PFK; UniProt ID: G3IDJ8), showing the overall enzyme structure. **(B)** Superposition of homologous PFK structures bound to phosphoenolpyruvate (PEP) and the structurally similar 2-phosphoglycolic acid (PGA) reveals that both ligands occupy equivalent inter-monomer binding sites within the dimer interface. **(C)** A zoomed-in view of the binding pocket indicates that, although PEP (modeled by Boltz-1) is positioned similarly to the homologous ligands, its carboxylic groups adopt the opposite orientation compared to those observed in Bacillus stearothermophilus PFK.

**Supplementary Figure 6.**
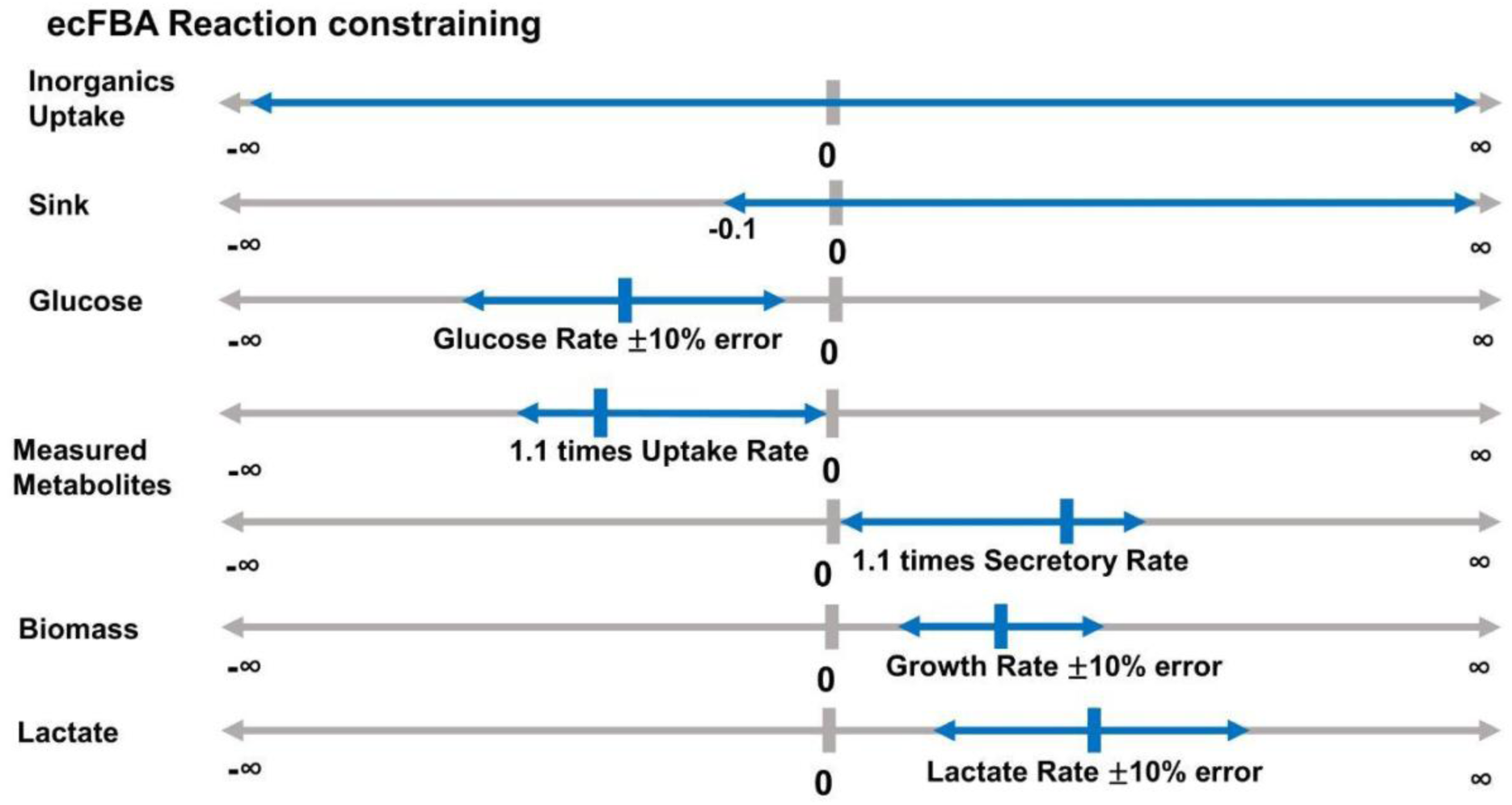
Schematic representation of the constraints applied in the ecFBA flux simulations: The diagram depicts how experimental measurement errors are integrated into the model by setting flux bounds. Specifically, glucose uptake, lactate secretion, and biomass production are each constrained to within ±10% of their measured values. In contrast, required inorganics are allowed free exchange (i.e., left unconstrained). All other measured metabolites, including amino acids, are constrained based on their direction of consumption/secretion. Secreted metabolites are restricted to flux values ranging from 0 up to the measured value+10% error margin, while uptake metabolites are limited to flux values from the measured value-10% up to 0. Additionally, sink reactions are constrained from –0.1 to infinity, permitting a small negative flux while allowing unlimited positive flux.

**Supplementary Table 7.** Whole cell concentrations of PEP and NADH in CHO-S wt and CHO-ZeLa strains during exponential growth: Absolute metabolomics samples were measured using LC-MS at 52.5 hours post inoculation in an ambr15 bioreactors. The measured concentrations were then converted to intracellular concentrations by using the VCD and average cell diameter. A two-sided Welch’s t-test was performed on the PEP concentrations with n = 8 samples and no t-test was possible for the NADH concentration as NADH was not detected (n.d.) in the CHO-S wt samples. Concentrations of the metabolites and their associated 95% confidence intervals are also presented. All statistical testing was performed on the log-transformed values before being converted into the mM concentrations given and with the exception of the NADH concentration that was only measured 3 times, both PEP concentrations were quantified with n = 4. Abbreviations -PEP: Phosphoenolpyruvate, NADH: Nicotinamide adenine dinucleotide (reduced).

| Met. | CHO-S wt (mM) (mean (95% CI)) | CHO-ZeLa (mM) (mean (95% CI)) |
| --- | --- | --- |
| PEP* (p = 0.03) | 3.3x10<sup>-3</sup> (2.9x10<sup>-4</sup> - 3.7x10<sup>-2</sup>) | 1.28x10<sup>-2</sup> (5.4x10<sup>-3</sup> - 3.1x10<sup>-2</sup>) |
| NADH | n.d. | 4.8x10<sup>-4</sup> (2.4x10<sup>-4</sup> - 9.3x10<sup>-4</sup>) |

## References

1. Tihanyi, B. & Nyitray, L. Recent advances in CHO cell line development for recombinant protein production. Drug Discov. Today Technol. 38, 25–34 (2020).

2. Kim, J. Y., Kim, Y.-G. & Lee, G. M. CHO cells in biotechnology for production of recombinant proteins: current state and further potential. Appl. Microbiol. Biotechnol. 93, 917–930 (2012).

3. Towards next generation CHO cell line development and engineering by systems approaches. Current Opinion in Chemical Engineering 22, 1–10 (2018).

4. Kyriakopoulos, S. et al. Kinetic Modeling of Mammalian Cell Culture Bioprocessing: The Quest to Advance Biomanufacturing. Biotechnol. J. 13, e1700229 (2018).

5. Ebrahimi, S. B. & Samanta, D. Engineering protein-based therapeutics through structural and chemical design. Nat. Commun. 14, 1–11 (2023).

6. Tao, F. et al. Digital twin-driven product design, manufacturing and service with big data. Int. J. Adv. Manuf. Technol. 94, 3563–3576 (2018).

7. Model-based assessment of mammalian cell metabolic functionalities using omics data. Cell Reports Methods 1, 100040 (2021).

8. Hefzi, H. et al. A Consensus Genome-scale Reconstruction of Chinese Hamster Ovary Cell Metabolism. Cell Syst 3, 434–443.e8 (2016).

9. Fouladiha, H. et al. Systematically gap-filling the genome-scale metabolic model of CHO cells. Biotechnol. Lett. 43, 73–87 (2021).

10. Yeo, H. C., Hong, J., Lakshmanan, M. & Lee, D.-Y. Enzyme capacity-based genome scale modelling of CHO cells. Metab. Eng. 60, 138–147 (2020).

11. Application of a curated genome-scale metabolic model of CHO DG44 to an industrial fed-batch process. Metabolic Engineering 51, 9–19 (2019).

12. Pang, K. T. et al. Genome-Scale Modeling of CHO Cells Unravel the Critical Role of Asparagine in Cell Culture Feed Media. Biotechnol J 19, e202400072 (2024).

13. Yusufi, F. N. K. et al. Mammalian Systems Biotechnology Reveals Global Cellular Adaptations in a Recombinant CHO Cell Line. Cell Syst 4, 530–542.e6 (2017).

14. Yeo, H. C. et al. Combined multivariate statistical and flux balance analyses uncover media bottlenecks to the growth and productivity of Chinese hamster ovary cell cultures. Biotechnol Bioeng 119, 1740–1754 (2022).

15. Hong, J. K. et al. Data-driven and model-guided systematic framework for media development in CHO cell culture. Metab Eng 73, 114–123 (2022).

16. Park, S.-Y. et al. Debottlenecking and reformulating feed media for improved CHO cell growth and titer by data-driven and model-guided analyses. Biotechnol J 18, e2300126 (2023).

17. Park, S.-Y. et al. Exploring metabolic effects of dipeptide feed media on CHO cell cultures by in silico model-guided flux analysis. Appl Microbiol Biotechnol 108, 123 (2024).

18. Gopalakrishnan, S. et al. Multi-omic characterization of antibody-producing CHO cell lines elucidates metabolic reprogramming and nutrient uptake bottlenecks. Metab Eng 85, 94– 104 (2024).

19. Gopalakrishnan, S. et al. COSMIC-dFBA: A novel multi-scale hybrid framework for bioprocess modeling. Metab Eng 82, 183–192 (2024).

20. Schinn, S.-M., Morrison, C., Wei, W., Zhang, L. & Lewis, N. E. A genome-scale metabolic network model and machine learning predict amino acid concentrations in Chinese Hamster Ovary cell cultures. Biotechnol Bioeng 118, 2118–2123 (2021).

21. Park, S.-Y. et al. Driving towards digital biomanufacturing by CHO genome-scale models. Trends Biotechnol 42, 1192–1203 (2024).

22. Gutierrez, J. M. et al. Genome-scale reconstructions of the mammalian secretory pathway predict metabolic costs and limitations of protein secretion. Nature Communications 11, 1– 10 (2020).

23. Strain, B., Morrissey, J., Antonakoudis, A. & Kontoravdi, C. How reliable are Chinese hamster ovary (CHO) cell genome-scale metabolic models? Biotechnol Bioeng 120, 2460– 2478 (2023).

24. Hefzi, H. et al. Multiplex genome editing eliminates lactate production without impacting growth rate in mammalian cells. Nature Metabolism 7, 212–227 (2025).

25. Brunk, E. et al. Recon3D enables a three-dimensional view of gene variation in human metabolism. Nat. Biotechnol. 36, 272–281 (2018).

26. Sayers, E. W. et al. Database resources of the National Center for Biotechnology Information. Nucleic Acids Res. 47, D23–D28 (2019).

27. UniProt Consortium, The. UniProt: the universal protein knowledgebase. Nucleic Acids Res. 46, 2699–2699 (2018).

28. Bansal, P. et al. Rhea, the reaction knowledgebase in 2022. Nucleic Acids Res. 50, D693– D700 (2022).

29. Kanehisa, M. & Goto, S. KEGG: kyoto encyclopedia of genes and genomes. Nucleic Acids Res. 28, 27–30 (2000).

30. Kim, S. et al. PubChem 2023 update. Nucleic Acids Res. 51, D1373–D1380 (2023).

31. King, Z. A. et al. BiGG Models: A platform for integrating, standardizing and sharing genome-scale models. Nucleic Acids Res. 44, D515 (2016).

32. Schellenberger, J., Park, J. O., Conrad, T. M. & Palsson, B. Ø. BiGG: a Biochemical Genetic and Genomic knowledgebase of large scale metabolic reconstructions. BMC Bioinformatics 11, 213 (2010).

33. Hastings, J. et al. ChEBI in 2016: Improved services and an expanding collection of metabolites. Nucleic Acids Res. 44, D1214–D1219 (2015).

34. Shlomi, T., Benyamini, T., Gottlieb, E., Sharan, R. & Ruppin, E. Genome-Scale Metabolic Modeling Elucidates the Role of Proliferative Adaptation in Causing the Warburg Effect. PLOS Computational Biology 7, e1002018 (2011).

35. Jumper, J. et al. Highly accurate protein structure prediction with AlphaFold. Nature 596, 583–589 (2021).

36. Wohlwend, J., et al. Boltz-1 Democratizing Biomolecular Interaction Modeling. bioRxiv (2024) doi:10.1101/2024.11.19.624167.

37. Varadi, M. et al. AlphaFold Protein Structure Database: massively expanding the structural coverage of protein-sequence space with high-accuracy models. Nucleic Acids Res. 50, D439–D444 (2022).

38. Robinson, J. L. et al. An atlas of human metabolism. Sci. Signal. 13, (2020).

39. Li, F., Chen, Y., Anton, M. & Nielsen, J. GotEnzymes: an extensive database of enzyme parameter predictions. Nucleic Acids Res. 51, D583–D586 (2022).

40. Xiong, K., et al. An optimized genome-wide, virus-free CRISPR screen for mammalian cells. Cell Reports Methods 1, (2021).

41. Bordbar, A., Monk, J. M., King, Z. A. & Palsson, B. O. Constraint-based models predict metabolic and associated cellular functions. Nature Reviews Genetics 15, 107–120 (2014).

42. Richelle, A., Chiang, A. W. T., Kuo, C.-C. & Lewis, N. E. Increasing consensus of context-specific metabolic models by integrating data-inferred cell functions. PLoS Comput Biol 15, e1006867 (2019).

43. Joshi, C. J. et al. StanDep: Capturing transcriptomic variability improves context-specific metabolic models. PLoS Comput. Biol. 16, e1007764 (2020).

44. Guidelines for extracting biologically relevant context-specific metabolic models using gene expression data. Metab. Eng. 75, 181–191 (2023).

45. Chandel, N. S. Amino Acid Metabolism. Cold Spring Harb Perspect Biol 13, (2021).

46. Lewis, N. E. et al. Omic data from evolved E. coli are consistent with computed optimal growth from genome-scale models. Mol Syst Biol 6, 390 (2010).

47. Zhang, Y. et al. Three-dimensional structural view of the central metabolic network of Thermotoga maritima. Science 325, 1544–1549 (2009).

48. Sánchez-Martínez, C. & Aragón, J. J. Analysis of phosphofructokinase subunits and isozymes in ascites tumor cells and its original tissue, murine mammary gland. FEBS Lett 409, 86–90 (1997).

49. Ozawa, K. Purification and kinetic properties of phosphofructokinase from dental pulps of rat incisors. Arch Oral Biol 30, 577–582 (1985).

50. Holness, M. J. & Sugden, M. C. Regulation of pyruvate dehydrogenase complex activity by reversible phosphorylation. Biochem Soc Trans 31, 1143–1151 (2003).

51. Patel, M. S. & Korotchkina, L. G. Regulation of the pyruvate dehydrogenase complex. Biochem Soc Trans 34, 217–222 (2006).

52. Mulukutla, B. C., Yongky, A., Daoutidis, P. & Hu, W.-S. Bistability in glycolysis pathway as a physiological switch in energy metabolism. PLoS One 9, e98756 (2014).

53. Berman, H. M. et al. The Protein Data Bank. Nucleic Acids Res 28, 235–242 (2000).

54. Mosser, R., Reddy, M. C. M., Bruning, J. B., Sacchettini, J. C. & Reinhart, G. D. Redefining the role of the quaternary shift in Bacillus stearothermophilus phosphofructokinase. Biochemistry 52, 5421–5429 (2013).

55. Thiele, I. & Palsson, B. Ø. A protocol for generating a high-quality genome-scale metabolic reconstruction. Nat Protoc 5, 93–121 (2010).

56. Marin de Mas, I., Herand, H., Carrasco, J., Nielsen, L. K. & Johansson, P. I. A Protocol for the Automatic Construction of Highly Curated Genome-Scale Models of Human Metabolism. Bioengineering (Basel) 10, (2023).

57. Ebrahim, A., Lerman, J. A., Palsson, B. O. & Hyduke, D. R. COBRApy: COnstraints-Based Reconstruction and Analysis for Python. BMC Syst. Biol. 7, 74 (2013).

58. Beguerisse-Díaz, M., Bosque, G., Oyarzún, D., Picó, J. & Barahona, M. Flux-dependent graphs for metabolic networks. NPJ Syst Biol Appl 4, 32 (2018).

59. Cock, P. J. A. et al. Biopython: freely available Python tools for computational molecular biology and bioinformatics. Bioinformatics 25, 1422–1423 (2009).

60. Yates, A. et al. The Ensembl REST API: Ensembl Data for Any Language. Bioinformatics 31, 143–145 (2015).

61. Széliová, D. et al. What CHO is made of: Variations in the biomass composition of Chinese hamster ovary cell lines. Metab. Eng. 61, 288–300 (2020).

62. Jeske, L., Placzek, S., Schomburg, I., Chang, A. & Schomburg, D. BRENDA in 2019: a European ELIXIR core data resource. Nucleic Acids Res. 47, D542–D549 (2019).

63. Wang, Y., Eddy, J. A. & Price, N. D. Reconstruction of genome-scale metabolic models for 126 human tissues using mCADRE. BMC Systems Biology 6, 1–16 (2012).

64. Chung, B. K. S. & Lee, D.-Y. Flux-sum analysis: a metabolite-centric approach for understanding the metabolic network. BMC Syst Biol 3, 117 (2009).

65. Heirendt, L. et al. Creation and analysis of biochemical constraint-based models using the COBRA Toolbox v.3.0. Nat Protoc 14, 639–702 (2019).

